# Reducing encapsidated impurity DNA derived from plasmid backbone by modifying the p5 terminal resolution site in rAAV vector production

**DOI:** 10.64898/2026.04.22.720036

**Authors:** Yuya Nishimura, Shoko Hataya, Shunsuke Saito, Naoki Makita

## Abstract

Recombinant adeno-associated virus (rAAV) vectors are pivotal for gene therapy; however, the encapsidation of residual DNA, particularly plasmid backbone sequences, pose significant safety risks. Recent studies have identified the p5 promoter, which contains a Rep-binding element and a terminal resolution site (TRS), as a cryptic origin of replication that facilitates packaging of upstream sequences. In this study, we investigated the effect of p5 TRS modifications on impurity DNA levels in a single-plasmid All-in-One (AiO) AAV production system. Wild-type p5 (p5wt) promoted significant packaging of upstream plasmid backbone DNA, especially when the backbone was positioned between p5wt and the inverted terminal repeat. Introducing mutations or deletions in the p5 TRS significantly reduced encapsidation of plasmid-derived sequences, including kanamycin resistance genes, and improved the ratio of full to partial particles, as seen with the p5Δloop variant. Furthermore, the p5Δloop-AiO system showed higher rAAV yields than both conventional triple-transfection methods and previously reported p5-spacer variants. Thus, our findings suggest a robust vector design strategy for minimizing DNA impurities, thereby enhancing the safety and efficacy of AAV-based gene therapy.

## Introduction

Adeno-associated virus (AAV) vectors, owing to their favorable safety profile and tissue tropism, are increasingly used as gene therapy modalities for various hereditary diseases. However, adverse events reported after high-dose systemic administration in clinical settings have renewed concerns about the risks associated with impaired vector product quality, particularly DNA impurities unintentionally introduced during manufacturing.^1,2,3^

While impurities outside AAV particles can be minimized through rigorous purification, nucleic acid impurities inside the particles are difficult to remove.^4^ Besides the target therapeutic expression cassette and its truncated sequences, two major types of nucleic acid impurities are packaged inside AAV particles.^5,6^ One consists of DNA fragments derived from the host genome of production cell lines such as HEK293 cells. The other comprises sequences derived from plasmids used during manufacturing, specifically cross-packaged plasmid backbone sequences located outside the inverted terminal repeats (ITRs).^7,8^ Furthermore, the generation of replication-competent adeno-associated virus (rcAAV), resulting from reconstitution of the *Rep* and *Cap* genes flanked by a pair of ITRs, has also been reported.^6,9^ These impurities may not only diminish therapeutic efficacy but also trigger unintended immune responses and genotoxicity.

Impurities derived from plasmid backbones pose particular safety concerns. Many manufacturing plasmids contain drug resistance genes as selection markers, and may persist in the human body through genomic integration or long-term expression.^10^ Moreover, the transcriptional activity of plasmid sequences near the ITR has been associated with neurotoxicity,^11^ indicating that backbone-derived DNA is not merely an inert impurity.

Recent studies have shown that sequences surrounding p5 are preferentially encapsidated.^12^ Although p5 is a promoter that regulates the expression of Rep78/68, it is also thought to function as a pseudo-packaging origin. This is because p5 contains a Rep-binding element and a terminal resolution site (TRS) similar to the ITR. Mispackaging of p5-derived DNA may increase the risk of rcAAV formation and contamination with impurities, including virus-derived genes such as *Rep* or *Cap* and plasmid backbone DNA.

Dual- and single-plasmid systems,^13,14^ in which ITR and p5 are located on the same plasmid, have been developed to streamline the AAV manufacturing workflow. However, regions sandwiched between these elements are particularly prone to impurity packaging. To overcome this issue, we analyzed impurity DNA near the p5 sequence of a single-plasmid system, in which the plasmid backbone lies between the 5′ ITR and p5. Our aim was to provide insights into improving vector safety for gene therapy through advanced design strategies that minimize DNA impurities.

## Results

### Removal of the p5 TRS reduced impurity DNA

To prevent nicking at the TRS, p5 mutants were constructed by introducing mutations into the stem–loop structure surrounding the TRS (Figure 1A). As shown in Figure 1B, recombinant adeno-associated virus (rAAV) was produced by triple transfection (TTF) using each pRC plasmid containing one of these p5 mutants. Encapsidated impurity DNA from sequences upstream of p5 was then compared (Figure 1C). We found that rAAV produced with wild-type p5 (p5wt) contained a large amount of the kanamycin resistance gene sequence (KmR) from the plasmid backbone located upstream. In contrast, impurity DNA was significantly reduced in rAAV samples generated with TRS-mutated p5 variants. When p5wt was positioned downstream of the *Cap* gene, impurity DNA derived from KmR also decreased; however, contamination originating from sequences upstream of *Cap* increased. These results indicate that p5wt promotes encapsidation of its upstream sequences regardless of its position on the plasmid.

**Figure 1.**
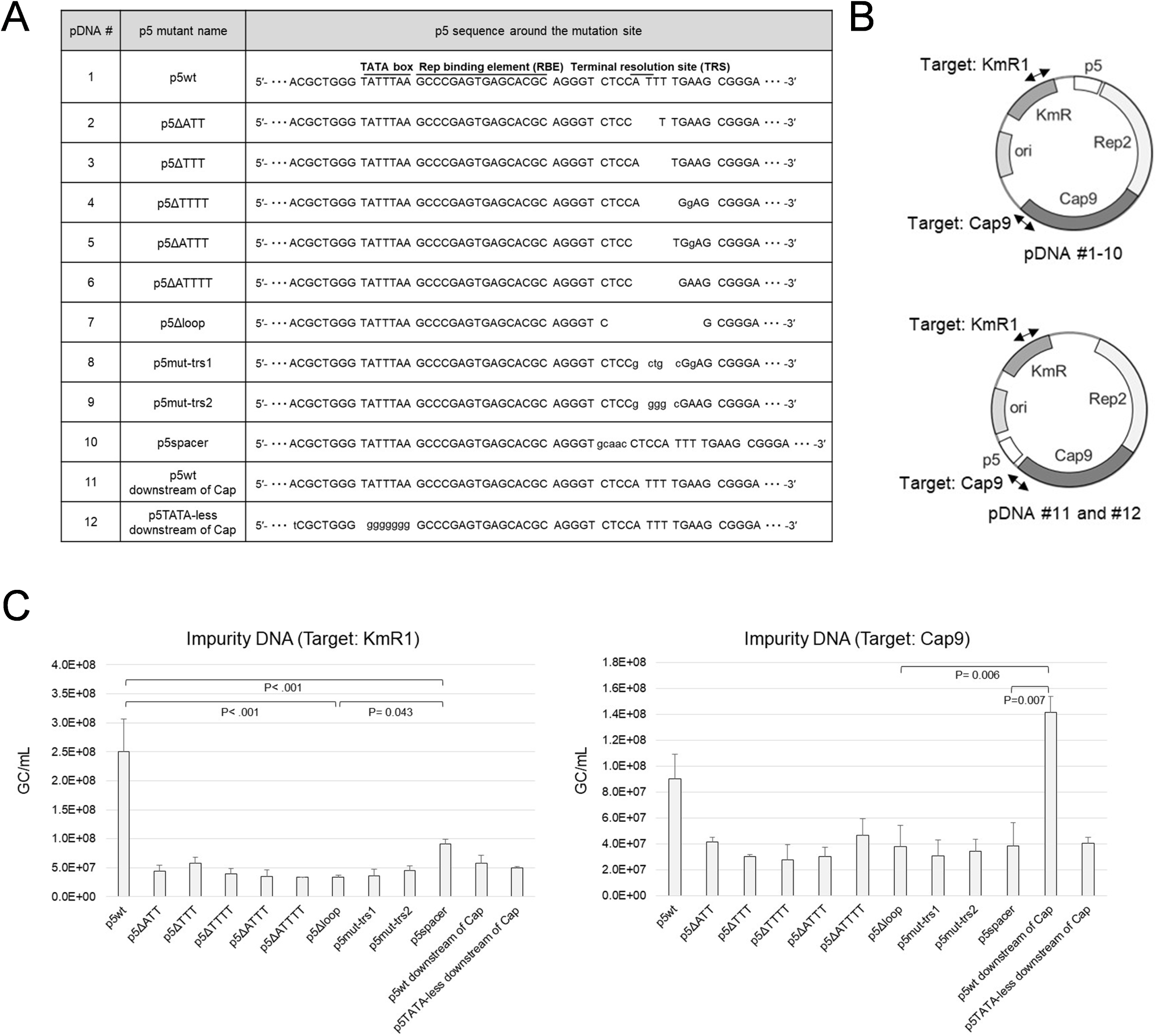
Evaluation of impurity DNA in recombinant adeno-associated virus (rAAV) vectors produced via triple transfection (TTF) using pRC constructs harboring diverse mutations in p5. (A) Names and sequences of each p5 mutant. (B) Schematic diagram showing gene arrangement in the pRC constructs and target positions for quantitative polymerase chain reaction (qPCR). (C) Quantification of impurity DNA derived from KmR1 and Cap9 sequences in rAAV. Viral Production Cells 2.0 were used, and experiments were carried out at the 6-well plate scale. *n* = 2 or 3; error bars indicate standard deviation (SD).

Furthermore, p5 mutants with deletions around the TRS or base substitutions that compromised the TRS sequence and stem-loop structure produced a greater reduction in impurity DNA than the previously reported p5spacer,^12^ which contains a 5-base spacer upstream of the TRS. Notably, rAAV produced with p5Δloop exhibited the lowest level of impurities derived from the plasmid backbone.

### Removal of the p5 TRS in All-in-One (AiO) constructs reduced plasmid backbone-derived impurity DNA

Single plasmids (AiO) incorporating the p5 mutants were constructed. To assess the effect of placing the ITR and p5 on the same plasmid, the plasmid backbone was positioned between p5 and the 5′ ITR (Figure 2A). Single transfection was used to produce rAAV, employing each AiO construct carrying a p5 mutant. The rAAV yield and levels of plasmid backbone-derived impurity DNA (kanamycin resistance gene sequence, KmR) packaged within the viral particles were then evaluated (Figure 2B).

**Figure 2.**
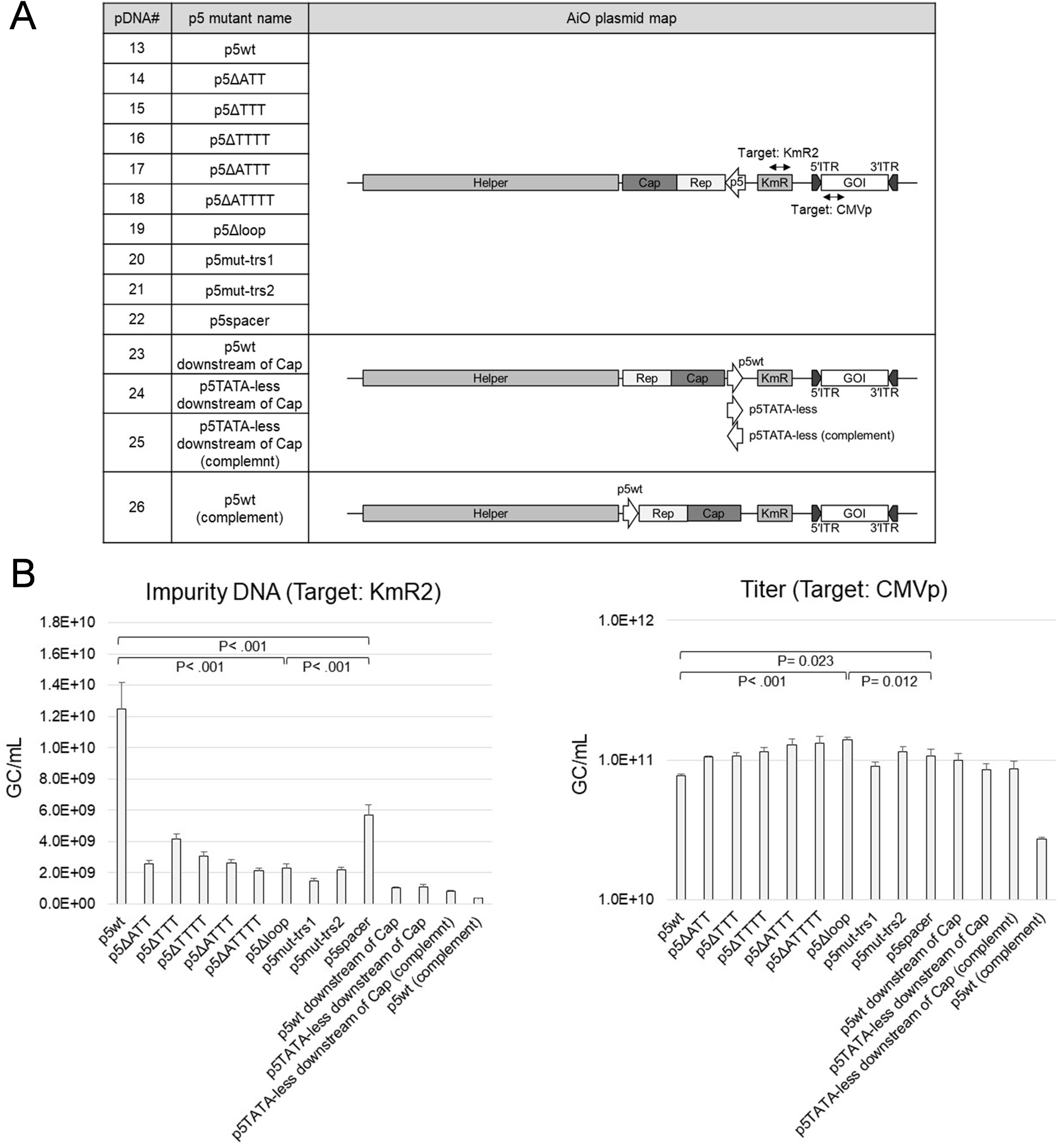
Evaluation of impurity DNA in rAAV produced via single transfection using all-in-one (AiO) plasmids harboring diverse mutations in p5. (A) Schematic diagram showing the arrangement, orientation, and qPCR target positions in the plasmids. (B) Quantification of impurity DNA (KmR2) and rAAV production yields. Viral Production Cells 2.0 were used, and the experiments were performed at the 6-well plate scale. *n* = 2 or 3; error bars indicate SD.

A substantial amount of impurity DNA was detected in AiO constructs possessing p5wt. However, reversing the orientation of p5 at the same position (p5wt downstream of *Cap*) significantly reduced impurity DNA levels. This finding suggests that when the ITR is located upstream of p5, the intervening plasmid backbone is readily encapsidated. However, even under this configuration, AiO constructs containing TRS-mutated p5 variants produced rAAV with markedly lower impurity DNA levels, similar to TTF.

Furthermore, AiO constructs carrying any of the p5 mutants exhibited lower impurity levels than those containing p5spacer. Notably, the AiO construct with p5Δloop also showed superior rAAV yield.

### p5 TRS had a greater impact on impurity DNA levels than the ITR in AiO

In rAAV produced using AiO constructs carrying each p5 mutant, impurity DNA was evaluated according to the distance from p5 (Figure 3). In AiO constructs containing p5wt, p5spacer, or p5ΔTTT, impurity DNA was clearly more abundant at +500 bp upstream of p5 than at +2500 bp (Figure 3A), which is closer to the 5′ ITR (approximately 700 bp from the 5′ ITR). Given that the TRS sequence is retained in p5 and its mutants, this finding indicates that nicking at the TRS strongly influences the generation of impurity DNA. Additionally, the higher impurity levels upstream of p5 than those upstream of the 5′ ITR suggest the possibility of ITR-independent packaging.

**Figure 3.**
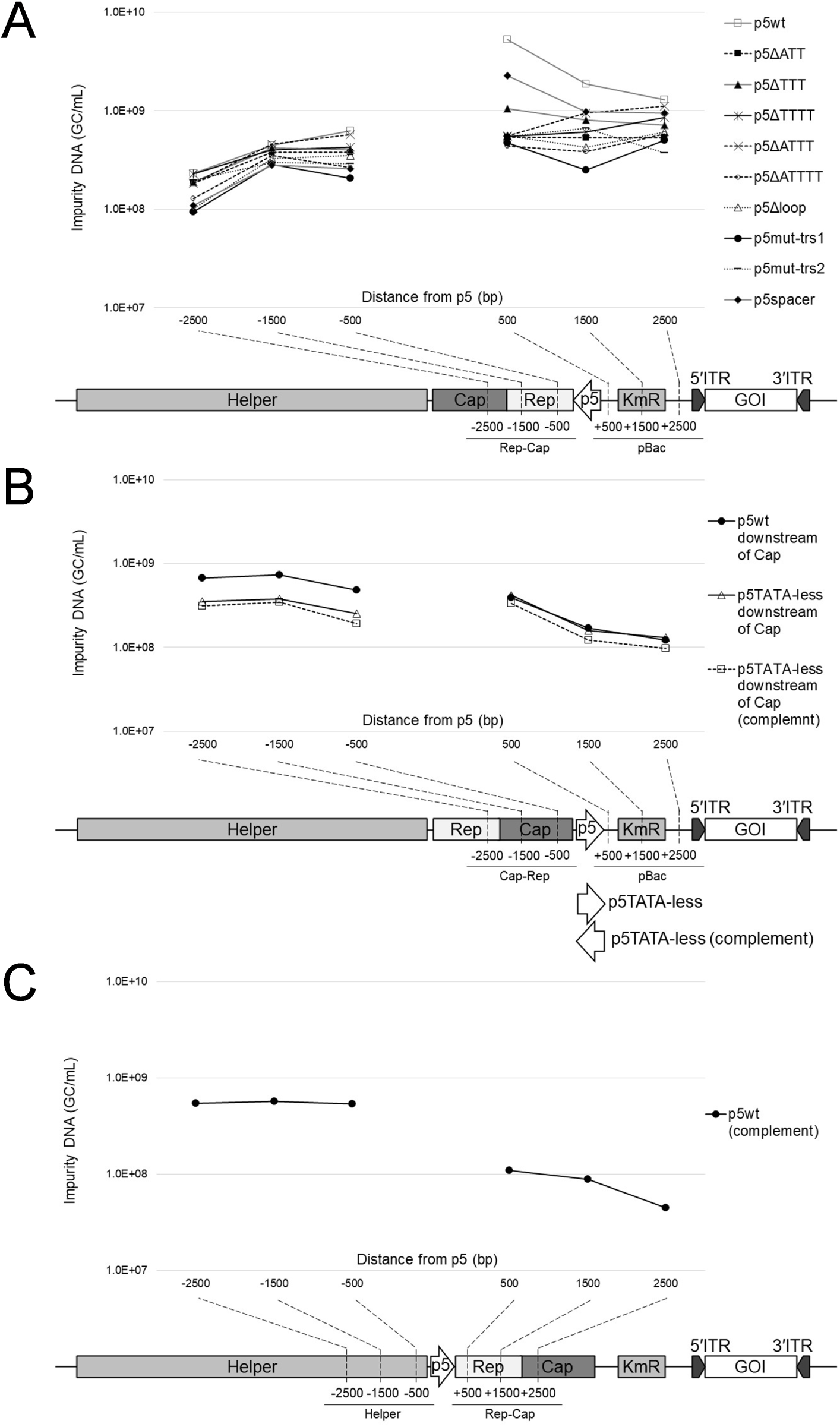
Evaluation of impurity DNA upstream and downstream of p5 mutants in rAAV produced via single transfection of AiO plasmids. Quantification of impurity DNA in regions upstream (+500, +1500, and +2500 bp) and downstream (-500 bp, -1500 bp, and -2500 bp) of p5, with the following gene arrangements: (A) pDNA#13-22, (B) pDNA#23-25, and (C) pDNA#26. Viral Production Cells 2.0 were used, and the experiments were conducted at the 6-well plate scale. *n* = 2 or 3.

As shown in Figure 3B, plasmid-derived impurity DNA decreased when p5wt orientation was reversed so that the 5′ ITR was no longer located upstream. However, sequences immediately upstream of p5 remained prone to impurity packaging. DNA impurities derived from the *Rep* and *Cap* genes (Figure 3B) and from the helper region (Figure 3C) were detected. In the p5TATA-less variant, which lacks promoter function, the upstream sequence was not prone to impurity packaging despite retention of the TRS sequence, and no orientation-related effect was observed.

### p5**Δ**loop reduced the production of partial particles

Genome integrity of packaged vectors was evaluated in rAAV produced via TTF using a pRC plasmid containing p5wt, via single transfection using an AiO construct containing p5wt, and via single transfection using an AiO construct containing p5Δloop. First, the ratio of particles containing partial genomes was estimated based on the difference in genome copy numbers between the ITR and the transgene *EGFP* (Figure 4A). Among the detected particles, 88.8, 89.8, and 65.5% were identified as partial particles for p5wt-TTF, p5wt-AiO, and p5Δloop-AiO, respectively.

**Figure 4.**
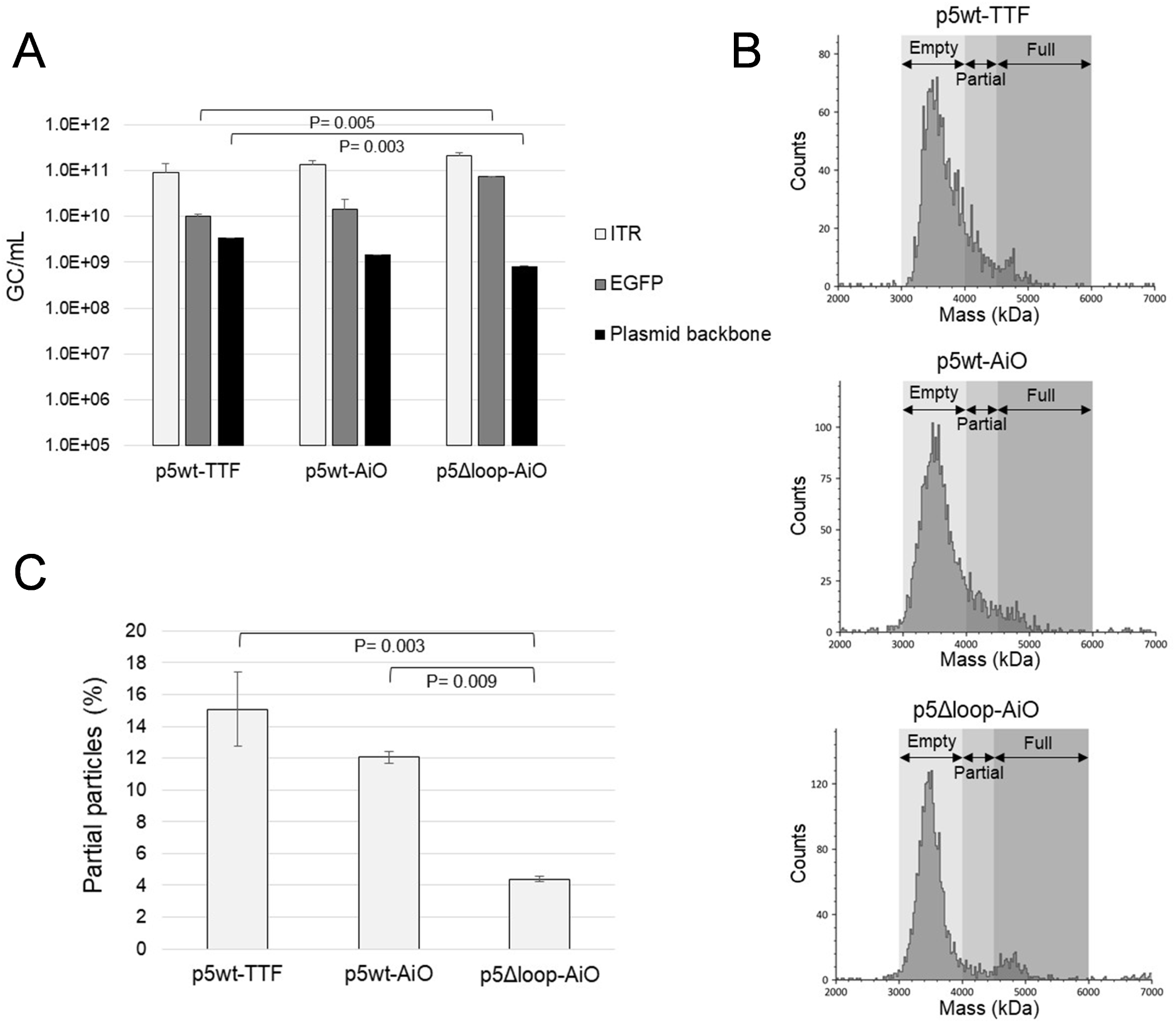
Comparison of impurity DNA and partial particles in rAAV produced using p5wt and p5Δloop constructs. rAAV carrying a 4 kb transgene was produced via TTF using pRC with p5wt, or via single transfection using AiO plasmids with p5wt or p5Δloop. Impurity DNA and partial particle content were compared. (A) Quantification using qPCR targeting ITR, *EGFP* (transgene), and the plasmid backbone. For p5wt-TTF, the plasmid backbone values represent the sum of AmpR from pRC and KmR2 from pGOI and the helper plasmid, which contains E2A, E4 and VA RNA. For p5wt-AiO or p5Δloop-AiO, plasmid backbone values represent only KmR2. (B) Mass photometry histograms showing molecular weight distribution. Particles of 3000–4000 kDa were classified as empty, 4000–4500 kDa as partial, and 4500–6000 kDa as full. (C) The ratio of partial particles was calculated from the total counts in (B). Suspension-adapted HEK293 cells were used as production cells, and experiments were conducted at the 125 mL flask scale. *n* = 2; error bars indicate SD.

Compared to full-like particles possessing the transgene (*EGFP*), the abundance ratios of particles carrying plasmid backbone sequences were 25.0, 9.4, and 1.1% for p5wt-TTF, p5wt-AiO, and p5Δloop-AiO, respectively. Thus, when p5wt was used, plasmid backbone-derive DNA impurities were higher in the rAAV produced using TTF than that produced using AiO constructs, although similar proportions of partial particles were observed in both. Additionally, replacing p5wt with a TRS-deleted p5 mutant, such as p5Δloop, reduced both partial particles and plasmid backbone-derived impurity DNA.

Next, the proportions of empty, partial, and full particles were quantified using mass photometry (Figure 4B). In rAAV produced using p5wt-TTF and p5wt-AiO, the empty peak exhibited tailing in the partial region. However, in rAAV produced using p5Δloop-AiO, clear separation was observed between the empty peak and full region. The abundance ratio of the particles is shown in Figure 4C. The proportions of partial particles in rAAV produced using p5wt-TTF, p5wt-AiO, and p5Δloop-AiO were 15.1, 12.1, and 4.4%, respectively, confirming that the p5Δloop mutation reduces the formation of partial particles.

## Discussion

Insertion of five bases upstream of the p5 TRS (mutant P5-HS5) reduces impurity DNA levels in AAV vectors. However, relatively high levels of impurities originating from the upstream sequences remain, as reported by Brimble et al. (2022).^12^ In the present study, although residual DNA was likewise reduced in P5-HS5 (notated as p5spacer) compared to that in p5wt, impurity levels remained higher than those observed in other mutants containing direct TRS mutations (Figure 1C, Figure 2B, Figure 3A). A similar pattern was observed for p5ΔTTT, a variant in which a continuous AT sequence remains at the TRS. These findings suggest that nicking at the TRS contributes substantially to the generation of impurity DNA.

A mutant in which the TATA box of p5 has been substituted with guanine residues (p5TATA-less) lacks promoter function for *Rep*, but is widely used downstream of *Cap* to maintain the production capacity of AAV.^15^ In the present study, impurity DNA near p5 was reduced in the p5TATA-less variant despite retention of the TRS, similar to the phenomenon observed in TRS-deficient mutants (Figure 2B, Figure 3B). This may be attributed to inefficient Rep-mediated cleavage at the TRS in the absence of TATA-binding protein, as noted by François et al. (2005).^16^

Even in a single-plasmid system, in which p5wt and the ITR were located on the same plasmid, plasmid backbone-associated impurities in rAAV produced via single transfection were lower than those in rAAV produced via TTF (Figure 4A). One possible explanation is that the majority of backbone impurities in TTF-derived rAAV originated from the gene-of-interest plasmid (pGOI), which contains ITRs. Consistently, in our study, impurities derived from the pGOI backbone accounted for 72.7% of all detected plasmid backbone sequences when p5wt was used (Supplemental Data 1).

Previous reports also indicate that impurity levels decrease with increasing plasmid backbone size.^17^ Accordingly, in the AiO system, plasmid backbone-associated impurities were likely reduced because the distance between the ITRs on the opposite side of the transgene exceeded the 4.7 kb packaging limit in AAV. Altogether, these findings suggest that ITR-derived impurity DNA, dominant in rAAV produced using TTF, decreases in AiO-derived vectors. Moreover, using AiO constructs with TRS-deleted p5 mutants may provide further reduction by minimizing p5-associated impurities.

In this study, only a limited variety of p5 mutants were tested within a single-plasmid system to focus on proof of concept, and the broader applicability of this methodology was not explored. Furthermore, evaluation of other impurity DNA species, such as rcAAV and host-derived contaminants, and their impact on the safety profile has not yet been carried out. Therefore, future studies should include conceptual validation using diverse cell lines, comprehensive mutagenesis or deletions in the vector design, full-sequence characterization of impurity DNA, and *in vivo* safety profiling and efficacy verification.

Overall, single transfection using AiO constructs containing p5 TRS mutants successfully reduced plasmid backbone-associated impurities encapsidated in rAAV. It increased vector yield and decreased the proportion of partial particles compared with conventional TTF. These findings suggest that AiO-based single transfection using TRS-modified p5 mutants may enable the production of rAAV with minimal DNA impurities, thereby improving vector quality for gene therapy applications.

## Materials and methods

### Plasmid construction

The origins of the vectors and inserts, as well as the primer numbers used for constructing the plasmids pDNA#1–#32, are provided in Supplemental Data 2. Primer sequences are listed in Supplemental Data 3. pDNA#1 was constructed by treating pUC19 (Addgene #500005, Watertown, MA, USA) with restriction enzymes AatII and PciI (New England Biolabs, Ipswich, MA, USA). Three polymerase chain reaction (PCR) fragments were inserted into the region from which the *LacZ* gene was removed. All fragment insertions were performed using In-Fusion Snap Assembly (Takara Bio Inc., Shiga, Japan). The p5 fragment was generated using primers #1 and #2 without template. The Rep2 fragment was amplified using primers #3 and #4 with pRC2-mi342 (Takara Bio Inc.) as the template. The SV40 *ori* fragment was amplified using primers #5 and #6 with pPB[Exp]-SV40>EBFP (VectorBuilder Inc., Chicago, IL, USA) as the template. The resulting plasmid (pR2) was treated with the restriction enzymes SwaI and AfeI, and a Cap9 fragment was inserted.

The Cap9 fragment was amplified using primers #7 and #8 with an artificial AAV9 *Cap* gene as template. The resulting plasmid (pDNA#30) was treated with restriction enzymes AsiSI and NotI, and three PCR fragments were inserted into the region from which the ampicillin resistance gene was removed. The SV40 *ori*-*ori* fragment was amplified using primers #9 and #10 with pR2 as template. The KmR fragment was amplified using primers #11 and #12 with pAAV[Exp]-CAG>EYFP:IRES:Neo (VectorBuilder) as template. The KmR promoter fragment was amplified using primers #13 and #14 with pR2 as template.

For pDNA#2–#9, pDNA#1 was treated with restriction enzymes NotI and SacI, and two PCR fragments were inserted into the region from which p5-Rep2 was removed. Primers #15 and #16 were used for the Rep2 region common to all constructs, primers #17 and #18 for the p5ΔATT fragment of pDNA#2, primers #17 and #19 for the p5ΔTTT fragment of pDNA#3, primers #17 and #20 for the p5ΔTTTT fragment of pDNA#4, primers #17 and #21 for the p5ΔATTT fragment of pDNA#5, primers #17 and #22 for the p5ΔATTTT fragment of pDNA#6, primers #17 and #23 for the p5Δloop fragment of pDNA#7, primers #17 and #24 for the p5mut-trs1 fragment of pDNA#8, and primers #17 and #25 for the p5mut-trs2 fragment of pDNA#9. pR2 was used as template for all aforementioned amplifications. For pDNA#10, pDNA#1 was treated with restriction enzymes NotI and SalI. Then, a PCR fragment amplified from pDNA#22 using primers #17 and #26 was inserted into the region from which p5-Rep2 was removed.

For pDNA#11 and #12, pDNA#1 was treated with restriction enzymes NotI and SacI, and a PCR fragment of FRT-Rep2 amplified from pRC2-mi342 (Takara) using primers #16 and #27 was inserted into the region from which p5-Rep2 was removed. The resulting plasmid was treated with restriction enzymes AsiSI and MfeI, and a PCR fragment amplified using primers #28 and #29 was inserted into the removed portion, using pR2 as the template for pDNA#11 and pRC2-mi342 (Takara) for pDNA#12. The plasmid SYNp177, which served as the backbone for pDNA#13, was synthesized by Synplogen Co., Ltd. (Kobe, Japan) using gene synthesis and the Ordered Gene Assembly in *Bacillus subtilis* (OGAB) method.^18,19^ Sequence information is provided in Supplemental Data 4. SYNp177 was treated with restriction enzymes FseI and SalI, and a PCR fragment amplified using primers #30 and #31 with pR2 as template was inserted at the site where the E1/p5/Rep2 fragments were removed. The resulting plasmid (SYNp180-1) was treated with restriction enzymes SwaI and AsiSI, and a Cap9 fragment amplified from pDNA#30 using primers #32 and #33 was inserted into the region from which the Cap1 gene was removed.

For pDNA#14–#21, pDNA#13 was treated with restriction enzymes NotI and SalI, and PCR-amplified p5 fragments were inserted into the region from which p5-Rep2 was removed. Primers #26 and #34 were used, with pDNA#2–#9 as templates.

For pDNA#22, pDNA#13 was treated with restriction enzymes NotI and SalI, and two PCR fragments were inserted into the region from which p5-Rep2 was removed. The p5spacer fragment was amplified using primers #35 and #36, while Rep2 fragment was amplified using primers #3 and #37. SYNp180-1 was used as template for both amplifications.

For pDNA#23, pDNA#13 was treated with restriction enzymes NotI and AsiSI, and a PCR fragment amplified from pDNA#11 using primers #38 and #39 was inserted into the region from which the p5-Rep2-Cap9 gene was removed.

For pDNA#24, pDNA#23 was treated with restriction enzymes PmeI and FseI, and a PCR fragment amplified from pDNA#12 using primers #39 and #40 was inserted into the region from which the p5 gene was removed.

For pDNA#25, pDNA#23 was treated with restriction enzymes PmeI and FseI, and a PCR fragment amplified from pDNA#12 using primers #41 and #42 was inserted into the region from which the p5 gene was removed.

For pDNA#26, SYNp180-1 was treated with restriction enzymes NotI and AsiSI, and a Rep2-Cap9 PCR fragment amplified from pDNA#13 using primers #43 and #44 was inserted into the region from which the Rep2-Cap1 gene was removed.

pDNA#27 (pAAV[Exp]-CMV>EGFP:WPRE) was purchased from VectorBuilder.

pDNA#28 was synthesized by Synplogen using gene synthesis and the OGAB method. Sequence information for this plasmid is provided in Supplemental Data 5.

For pDNA#29, pAAV-CMV (Takara) was treated with restriction enzymes AflII and BamHI, and four fragments were inserted into the removed portion. The eGFP fragment was amplified using primers #45 and #46 with pDNA#27 as template; the IRES fragment was amplified using primers #47 and #48 with pAAV[Exp]-CAG>EYFP:IRES:Neo (VectorBuilder) as template; the Hygromycin resistance gene fragment was amplified using primers #49 and #50 with pAAV[Exp]-EF1A>dTomato(ns):T2A:Hygro (VectorBuilder) as template; and the WPRE fragment was amplified using primers #51 and #52 with pDNA#27 as template.

The resulting plasmid (SYNp1163) was treated with restriction enzymes AseI and KasI, and a PCR fragment amplified from the gene-synthesized KmR-*ori* gene using primers #53 and #54 was inserted into the region from which the AmpR-*ori* gene was removed. The KmR-*ori* gene was synthesized by Synplogen using gene synthesis and the OGAB method, and the sequence information is provided in Supplemental Data 6.

For pDNA#31 and #32, pDNA#13 and #19 were treated with restriction enzymes NdeI and PacI, and a PCR fragment amplified from pDNA#29 using primers #52 and #55 was inserted into the region from which the eGFP-WPRE fragment was removed.

### Cell culture

Viral Production Cells 2.0 (VPC 2.0; Thermo Fisher Scientific, Waltham, MA, USA), a suspension cell line, were used. Cells were cultured in 125 mL plain-bottom Erlenmeyer flasks (125F) using Viral Production Medium (Gibco; Thermo Fisher Scientific) supplemented with 4 mM GlutaMAX (Gibco) and 1% penicillin–streptomycin (Nacalai Tesque, Inc., Kyoto, Japan).

HEK293 cells (ATCC, Manassas, VA, USA), adapted for suspension culture, were cultured in 125 mL baffled-bottom Erlenmeyer flasks (125F) using Cellvento 4HEK medium (Merck KGaA, Darmstadt, Germany) supplemented with 6 mM GlutaMAX (Gibco).

For culture and agitation, a New Brunswick S41i CO□ incubator shaker (Eppendorf, Hamburg, Germany) was used, with conditions set at 37□, 8% CO□, and 125 rpm. When the cell density reached 4–6 × 10□ cells/mL, subculturing was performed every 3 to 4 days using fresh medium to reach a density of 0.6 × 10□ cells/mL.

### Transfection

On the day of transfection, VPC 2.0 cells were seeded at a density of 3.0 × 10□ cells/mL (1.5 mL/well) in non-adhesive 6-well plates. For rAAV production, TransIT-VirusGen Select Transfection Reagent (Mirus Bio, Madison, WI, USA) was used. Plasmid DNA was added at 1.0 µg/10□ cells, and the transfection reagent was mixed with DNA at a ratio of 1.5:1 (v/w) to form complexes in fresh culture medium. The total volume of the complex solution was adjusted to 10% of the initial culture volume. After 30 min, the solution was added directly to the cell suspension in the 6-well plates.

HEK293 cells were seeded at 2.0 × 10□ cells/mL (30 mL/125F flask) on the day of transfection. FectoVIR-AAV (Polyplus-Transfection, Illkirch, France) was used as the transfection reagent for rAAV production. Plasmid DNA was added at 1.0 µg/10□ cells, and the transfection reagent was mixed with DNA at a ratio of 1:1 (v/w) to form complexes in fresh culture medium. The volume of the complex solution was adjusted to 5% of the cell culture volume. After 30 min, RevIT AAV Enhancer (Mirus Bio) was added to the complex solution at 0.1% of the initial culture volume, followed by direct addition to the cell suspension.

### Cell harvesting and lysis

Three days after transfection, cells were lysed by adding 20× lysis buffer (10% Tween 20 [Promega, Madison, WI, USA], 40 mM magnesium chloride solution [SERVA Electrophoresis GmbH, Heidelberg, Germany], and 2,000 units/mL Benzonase Nuclease HC [Novagen; Merck KGaA, Darmstadt, Germany]) to the cell culture medium to obtain a final buffer concentration of 1×. The mixture was incubated at 37□ for 1 h with agitation and cell debris was removed via centrifugation (12,000 g, 24□, 10 min). Poloxamer 188 (Sigma-Aldrich, St. Louis, MO, USA) was added to the supernatant (containing cell extract) to a final concentration of 0.001%. This was done to prevent the adsorption of proteins and viruses. The resulting lysate was used for downstream analyses.

### Quantification of DNA encapsulated in rAAV

To quantify rAAV genome titers, samples were treated with DNase I (Thermo Fisher Scientific, Vilnius, Lithuania) at 37□ for 30 min to remove residual plasmid DNA. Subsequently, EDTA (0.5 mol/L, diluted 100-fold; Nacalai Tesque, Inc.) was added, and samples were heated at 95□ for 10 min to stop the enzymatic reaction and release the encapsidated genome. Quantitative PCR (qPCR) was performed using a 100-fold dilution of the treated sample as template and Power SYBR Green PCR Master Mix (Applied Biosystems, Thermo Fisher Scientific, Austin, TX, USA). Amplification was carried out on a QuantStudio 3 Real-Time PCR System (Applied Biosystems; Thermo Fisher Scientific), with an initial incubation at 95□ for 10 s, followed by 40 cycles at 95□ for 10 s, 55□ for 5 s, and 72□ for 30 s.

For absolute quantification, plasmid DNA containing the target sequences was linearized using restriction enzymes, and serially diluted to generate a series of known copy number concentrations for use as a standard curve. The following target and primer combinations were used:

- KmR1 (primers #56 and #57) and Cap9 (primers #58 and #59), as shown in Figure 1.
- KmR2 (Primer #60 and #61) and CMVp (primers #62 and #63), as shown in Figure 2.
- Rep-Cap (+/-2500) (primers #64 and #65), Rep-Cap (+/-1500) (primers #66 and #67), Rep-Cap (+/-500) (primers #68 and #69), pBac (+500) (primers #70 and #71), pBac (+1500) (primers #72 and #73), pBac (+2500) (primers #74 and #75), Cap-Rep (-2500) (primers #76 and #77), Cap-Rep (-1500) (primers #78 and #79), Cap-Rep (-500) (primers #80 and #81), Helper (-2500) (primers #82 and #83), Helper (-1500) (primers #84 and #85), and Helper (-500) (primers #86 and #87), as shown in Figure 3.
- ITR (primers #88 and #89), EGFP (primers #90 and #91), AmpR (primers #92 and #93), and KmR2 (primers #60 and #61), as shown in Figure 4.

Subsequently, the absolute amount was calculated based on the standard curve for each target. To correct for errors due to inter-plate variation and differences in primer sets, a common control sample was included across all plates and used for normalization to ensure comparison validity between samples.

### Evaluation of genome packaging using mass photometry

To recover rAAV particles from the lysed sample liquid, KingFisher Duo Prime (Thermo Fisher Scientific) and Dynabeads CaptureSelect AAVX (Thermo Fisher Scientific) were used. Samples, magnetic beads, washing buffer (D-PBS, Gibco), and elution buffer [0.1 M glycine-HCl (pH 2.5)] (Nacalai Tesque, Inc.) were dispensed into 96-deep-well plates and set in the device according to the manufacturer’s protocol.

The magnetic beads and samples were mixed together, and shaken at room temperature (24□) for 30 min to specifically bind AAV9 particles. After washing to remove non-specifically adsorbed components, the bound AAV9 particles were recovered using an elution buffer. The eluate pH was immediately adjusted using a neutralization buffer containing 500 mM bis-tris propane (Tokyo Chemical Industry Co., Ltd., Tokyo, Japan), 750 mM NaCl (Nacalai Tesque, Inc.), and 0.05% Poloxamer 188 (pH 10.0; Sigma-Aldrich).

Molecular weight distribution of the particles was evaluated using a Refeyn Samux MP mass photometer (Refeyn Ltd., Oxford, UK). The samples were dropped onto cover glasses using silicon gaskets, and videos were recorded for 1 min. Data were analyzed using DiscoverMP software (Refeyn Ltd., Oxford, UK). Calibration was performed using a 660 kDa thyroglobulin standard solution (MARUWA Pharma Bio-Tech, Aichi, Japan), and molecular weight distribution was calculated from the resulting calibration curve.

### Statistical analysis

For the data shown in Figures 1 and 2, one-way analysis of variance was performed, followed by Tukey’s post hoc test in Python for multiple group comparisons. For the data shown in Figure 4, t-tests were performed using Microsoft Excel to compare two groups. A *p*-value < 0.05 was considered statistically significant.

## Data availability

The data supporting this study are available from the corresponding author upon reasonable request.

## Acknowledgments

The authors wish to thank Dr. Kenji Tsuge for his assistance with the synthesis of plasmid DNA and Editage (www.editage.jp) for English language editing. This study received no external funding.

## Author contributions

Conceptualization: YN, Data curation: YN, Formal analysis: YN, Investigation: YN, SH, Methodology: YN, Project administration: NM, SS, Supervision: NM, SS, Validation: YN, Visualization: YN, Writing–original draft: YN, Writing–review & editing: YN, SH, NM, SS

