## Supplementary Information for "Reducing encapsidated impurity DNA derived from plasmid backbone by modifying the p5 terminal resolution site in rAAV vector production"

Supplemental data 1

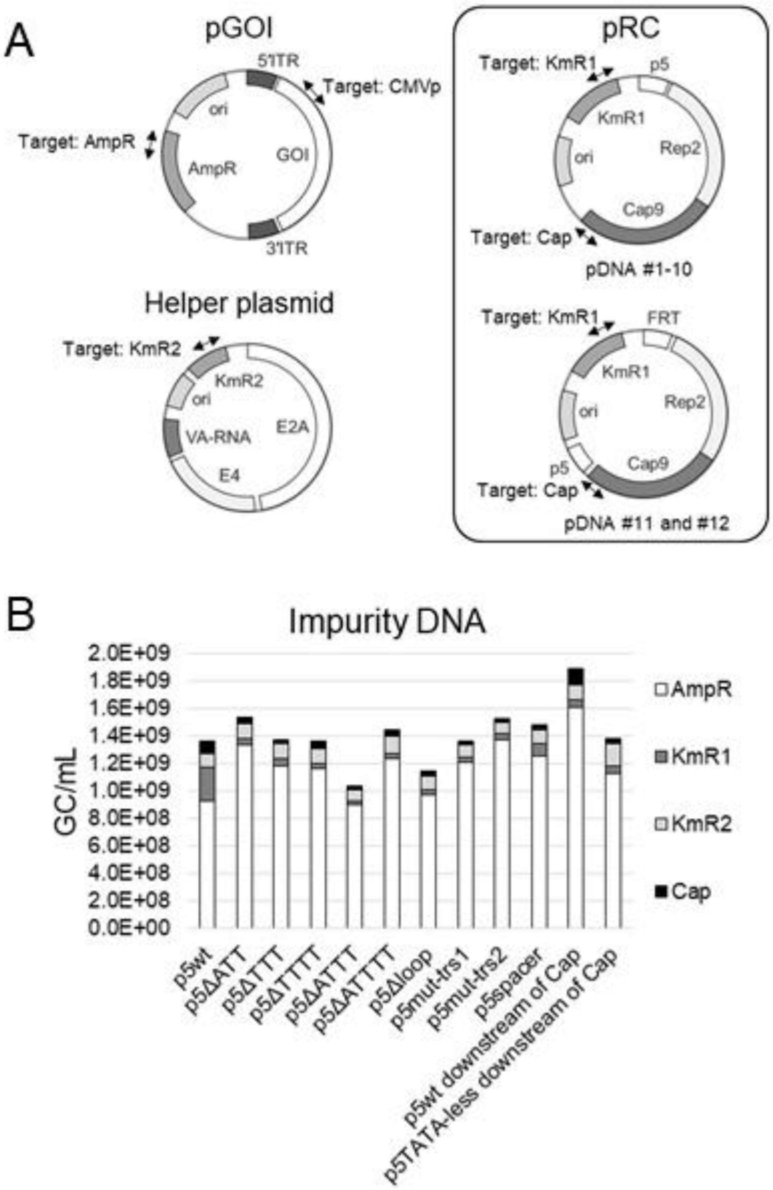


Supplemental Figure 1. Evaluation of total impurity DNA derived from plasmid backbone in rAAV vectors produced via TTF using pRC constructs harboring diverse mutations in p5 (A) Schematic diagram showing gene arrangement in the pGOI, helper plasmid and pRC constructs and target positions for qPCR. (B) Quantification of impurity DNA derived from AmpR, KmR1, KmR2 and Cap9 sequences in rAAV. Viral Production Cells 2.0 were used, and experiments were carried out at the 6-well plate scale. n = 2 or 3; error bars indicate SD.

Supplemental data 2

Table 1. List of vectors, templates and primers used for plasmid construction

| pDNA# | Plasmid ID | Vector | Template sequence | Primer # |
|---|---|---|---|---|
| 1 | SYNp69 | pUC19 (Addgene: #500005) | - | 1 and 2 |
| | | | pRC2-mi342 (Takara) | 3 and 4 |
| | | | pPB[Exp]-SV40>EBFP (VectorBuilder) | 5 and 6 |
| | | pR2 (This study) | GenBank: AY530579.1 | 7 and 8 |
| | | SYNp29 (This study) | pR2 (This study) | 9 and 10 |
| | | | pAAV[Exp]-CAG>EYFP:IRES:Neo (VectorBuilder) | 11 and 12 |
| | | | pR2 | 13 and 14 |
| 2 | SYNp1370 | SYNp69 (This study) | pR2 | 15 and 16 |
| | | | pR2 | 17 and 18 |
| 3 | SYNp1371 | SYNp69 | pR2 | 15 and 16 |
| | | | pR2 | 17 and 19 |
| 4 | SYNp1372 | SYNp69 | pR2 | 15 and 16 |
| | | | pR2 | 17 and 20 |
| 5 | SYNp1373 | SYNp69 | pR2 | 15 and 16 |
| | | | pR2 | 17 and 21 |
| 6 | SYNp482 | SYNp69 | pR2 | 15 and 16 |
| | | | pR2 | 17 and 22 |
| 7 | SYNp485 | SYNp69 | pR2 | 15 and 16 |
| | | | pR2 | 17 and 23 |
| 8 | SYNp483 | SYNp69 | pR2 | 15 and 16 |
| | | | pR2 | 17 and 24 |
| 9 | SYNp484 | SYNp69 | pR2 | 15 and 16 |
| | | | pR2 | 17 and 25 |
| 10 | SYNp1369 | SYNp69 | SYNp180-9-2 (This study) | 17 and 26 |
| 11 | SYNp481 | SYNp69 | pRC2-mi342 | 16 and 27 |
| | | | pR2 | 28 and29 |
| 12 | SYNp480 | SYNp69 | pRC2-mi342 | 16 and 27 |
| | | | pRC2-mi342 | 28 and29 |
| 13 | SYNp180-9 | SYNp177 (This study, Supplemental data 3) | pR2 | 30 and 31 |
| | | SYNp180-1 (This study) | SYNp29 | 32 and 33 |
| 14 | SYNp1404 | SYNp180-9 (This study) | SYNp1370 (This study) | 26 and 34 |
| 15 | SYNp1405 | SYNp180-9 | SYNp1371 (This study) | 26 and 34 |
| 16 | SYNp1406 | SYNp180-9 | SYNp1372 (This study) | 26 and 34 |
| 17 | SYNp1407 | SYNp180-9 | SYNp1373 (This study) | 26 and 34 |
| 18 | SYNp510 | SYNp180-9 | SYNp482 (This study) | 26 and 34 |
| 19 | SYNp513 | SYNp180-9 | SYNp485 (This study) | 26 and 34 |
| 20 | SYNp511 | SYNp180-9 | SYNp483 (This study) | 26 and 34 |
| 21 | SYNp512 | SYNp180-9 | SYNp484 (This study) | 26 and 34 |
| 22 | SYNp180-9-2 | SYNp180-9 | SYNp180-1 | 35 and 36 |
| | | | SYNp180-1 | 3 and 37 |
| 23 | SYNp487 | SYNp180-9 | SYNp481 (This study) | 38 and 39 |
| 24 | SYNp488 | SYNp487 (This study) | SYNp480 (This study) | 39 and40 |

| | | | | |
|---|---|---|---|---|
| 25 | SYNp489 | SYNp487 | SYNp480 | 41 and 42 |
| 26 | SYNp180-9-1 | SYNp180-1 | SYNp180-9 | 43 and 44 |
| 27 | pAAV[Exp]-CMV>EGFP:WPRE (VectorBuilder) | - | - | - |
| 28 | SYNp1088 (Supplemental data 4) | - | - | - |
| 29 | SYNp1164 | pAAV-CMV (Takara) | pAAV[Exp]-CMV>EGFP:WPRE (VectorBuilder) | 45 and 46 |
| | | | pAAV[Exp]-CAG>EYFP:IRES:Neo (VectorBuilder) | 47 and 48 |
| | | | pAAV[Exp]-EF1A>dTomato(ns):T2A:Hygro (VectorBuilder) | 49 and 50 |
| | | | pAAV[Exp]-CMV>EGFP:WPRE (VectorBuilder) | 51 and 52 |
| | | SYNp1163 (This study) | KmR-ori (Supplemental data 5) | 53 and 54 |
| 30 | SYNp29 | pR2 | GenBank: AY530579.1 | 7 and 8 |
| 31 | SYNp1198 | SYNp180-9 | SYNp1164 (This study) | 52 and 55 |
| 32 | SYNp1199 | SYNp513 | SYNp1164 | 52 and 55 |

Supplemental data 3

Table 2. List of primer sequences

| Primer # | Sequence (5′-3′) |
|---|---|
| 1 | gaaaagtgccacctgacgtcgcggccgcggaggggtggagtcgtgacgtgaattacgtcatagggttaggga ggtcctgtattagaggtcacgtgagtgttttgcgac |
| 2 | gttcaaacctcccgcttcaaaatggagaccctgcgtgctcactcgggcttaaatacccagcgtgaccacatggt gtcgcaaaatgtcgcaaaacactca |
| 3 | gcgggaggtttgaacgcgcagccgccatgccggggttttac |
| 4 | actagtagcgctatttaaatcatttattgttcaaagat |
| 5 | aatagcgctgcgatcgcggcctccaaaaaagc |
| 6 | ccttttgctcacatgtcaattgatcccgcccctaact |
| 7 | aacaataaatgatttaaatcagg |
| 8 | ggccgcgatcgcagcgctgtagccatggaaactagata |
| 9 | ggctacagcgctgcg |
| 10 | gttcttctgactgtcagaccaagtttactc |
| 11 | gagtaaacttggtctgacagtcagaagaactcgtc |
| 12 | aatattgaaaaaggaagagtatgattgaacaagat |
| 13 | gttcaatcatactcttcctttttcaa |
| 14 | tcacgactccacccctcc |
| 15 | cgggaggtttgaacgcgcagc |
| 16 | gtccacgcccactggagctcaggctgggttttggg |
| 17 | cctgacgtcgcggccgcggaggggtggagtcgtgacg |
| 18 | gctgcgcgttcaaacctcccgcttcaaggagaccctgcgtgctcactc |
| 19 | gcatggcggctgcgcgttcaaacctcccgcttcatggagaccctgcgtgctcac |
| 20 | gcatggcggctgcgcgttcaaacctcccgctcctggagaccctgcgtgctcac |
| 21 | gcatggcggctgcgcgttcaaacctcccgctccaggagaccctgcgtgctcactc |
| 22 | ctgcgcgttcaaacctcccgcttcggagaccctgcgtgctcactcgg |
| 23 | ctgcgcgttcaaacctcccgcgaccctgcgtgctcactcgg |
| 24 | ctgcgcgttcaaacctcccgctccgcagcggagaccctgcgtgctcactcgg |
| 25 | ctgcgcgttcaaacctcccgcttcgccccggagaccctgcgtgctcactcgg |
| 26 | ccagatcaccatcttgtcgacacag |
| 27 | cctgacgtcgcggccgccaggaaacagctatgacc |
| 28 | tatctagtttccatggctacgtagataagtagcatggcgggttaatcattaactacagcccgggcgtttaaacag cgggcggaggggtggagtcgtgacg |
| 29 | gctcacatgtcaattggttcaaacctcccgcttcaa |
| 30 | tcaccatcttgtcgacacagtcgttgaaggg |

| | |
|---|---|
| 31 | agcgcagcgagtcggccggccgcggccgcggaggggtggagtcg |
| 32 | gtgagcaaaagggcgatcgctctagaactagtgtagccatggaaactagata |
| 33 | aacaataaatgatttaaatcaggtatggctgccga |
| 34 | cgagtcggccggccgcggccgcggaggggtggagtcgtgacg |
| 35 | cggccggccgcggccgcggaggggtggagtcgtgacg |
| 36 | gttcaaacctcccgcttcaaaatggaggttgcaccctgcgtgctcactcggg |
| 37 | tcaccatcttgtcgacacag |
| 38 | gacatgtgagcaaaagggcgatcgccaggaaacagctatgaccatg |
| 39 | gaccgagcgcagcgagtcggccggccgttcaaacctcccgcttcaa |
| 40 | taactacagcccgggcgtttaaacagcgggcggaggggtg |
| 41 | taactacagcccgggcgtttaaacagcgggcgttcaaacctcccgcttcaa |
| 42 | gaccgagcgcagcgagtcggccggccggaggggtggagtcgtgacg |
| 43 | gtgagcaaaagggcgatcgcggaggggtggagtcgtgacg |
| 44 | cggccggccgcggccgctctagaactagtgtagccatggaaac |
| 45 | taagcagagctggtttagtg |
| 46 | gagggagaggggcgaattcttacttgtacagctcgtccatg |
| 47 | atggacgagctgtacaagtaagaattcgcccctctccctcccccccc |
| 48 | tcaggctttttcatgtcgacggttgtggccatattatcatc |
| 49 | accgtcgacatgaaaaagcctgaactcac |
| 50 | gttgattctcgagctattcctttgccctcggac |
| 51 | ggaatagctcgagaatcaacctctggattacaa |
| 52 | gggtcacagggatgccacccggatccgcggggaggcggcccaaagg |
| 53 | gcgcagcctgaatggcgaatggccggccatttgcggccgc |
| 54 | gagtgagctaactcacattaatgggattttgccgatttcggc |
| 55 | agtacatcaagtgtatcatatgcc |
| 56 | aacaagatggattgcacgcagg |
| 57 | cgattgtctgttgtgcccagtc |
| 58 | ccaaaattcctcacacggac |
| 59 | caggtacaggtgtgtttttg |
| 60 | atgattgaacaggatggcctgc |
| 61 | ctgcgcccaatcatagccaaac |
| 62 | catcaatgggcgtggatagc |
| 63 | ggagttgttacgacattttggaaa |
| 64 | ccacctgaagccattgtaag |
| 65 | ctcaatttcggtcagactgg |
| 66 | ggctcacttatatctgcgtc |

| | |
|---|---|
| 67 | caaaggatcacgtggttgag |
| 68 | gttttggggagcaagtaattg |
| 69 | ggagttgttacgacattttggaaa |
| 70 | cgccttatccggtaactatc |
| 71 | catacctcgctctgctaatc |
| 72 | ctagacagcagatcctgacc |
| 73 | gaacgagctgcaggatgaag |
| 74 | ggaagagttcttgcagctcg |
| 75 | caccatgaccgagtacaagc |
| 76 | gcgtatcagaaactgtgctac |
| 77 | catccaaatccacattgacc |
| 78 | cagattccactgccacttctc |
| 79 | gaagagcttgaagttgagtcg |
| 80 | ccaaaattcctcacacggac |
| 81 | caggtacaggtgtgtttttg |
| 82 | ggagcgcgttcacttaatag |
| 83 | gaagagctggaagaaccatg |
| 84 | gatttaagtgagatcagggtgc |
| 85 | ggtctcctcagcgatgattc |
| 86 | cttggaaaataaccctccgg |
| 87 | ctagctttttggccactgg |
| 88 | ggaacccctagtgatggagtt |
| 89 | cggcctcagtgagcga |
| 90 | gtccgccctgagcaaaga |
| 91 | tccagcaggaccatgtgatc |
| 92 | tcaacatttccgtgtcgccc |
| 93 | ttcaccagcgtttctgggtg |

Supplemental data 4

Sequence of SYNp177

cctcgaggcctgcaggcagctgcgcgctcgctcgctcactgaggccgcccgggcaaagcccgggcgtcgggcgacctttggtcgcc
cggcctcagtgagcgagcgagcgcgcagagagggagtggccaactccatcactaggggttcctatcgatatcaagctttaatagtaa
tcaattacggggtcattagttcatagcccatatatggagttccgcgttacataacttacggtaaatggcccgcctggctgaccgcccaa
cgacccccgcccattgacgtcaataatgacgtatgttcccatagtaacgccaatagggactttccattgacgtcaatgggtggagtattt
acggtaaactgcccacttggcagtacatcaagtgtatcatatgccaagtacgccccctattgacgtcaatgacggtaaatggcccgcct
ggcattatgcccagtacatgaccttatgggactttcctacttggcagtacatctacgtattagtcatcgctattaccatggtgatgcggtt
ttggcagtacatcaatgggcgtggatagcggtttgactcacggggatttccaagtctccaccccattgacgtcaatgggagtttgttttg
gcaccaaaatcaacgggactttccaaaatgtcgtaacaactccgccccattgacgcaaatgggcggtaggcgtgtacggtgggaggt
ctatataagcagagctggtttagtggatatcgtatacgccaccatggtgagcaagggcgaggagctgttcaccggggtggtgcccatc
ctggtcgagctggacggcgacgtaaacggccacaagttcagcgtgtccggcgagggcgagggcgatgccacctacggcaagctga
ccctgaagttcatctgcaccaccggcaagctgcccgtgccctggcccaccctcgtgaccaccctgacctacggcgtgcagtgcttcag
ccgctaccccgaccacatgaagcagcacgacttcttcaagtccgccatgcccgaaggctacgtccaggagcgcaccatcttcttcaag
gacgacggcaactacaagacccgcgccgaggtgaagttcgagggcgacaccctggtgaaccgcatcgagctgaagggcatcgact
tcaaggaggacggcaacatcctggggcacaagctggagtacaactacaacagccacaacgtctatatcatggccgacaagcagaag
aacggcatcaaggtgaacttcaagatccgccacaacatcgaggacggcagcgtgcagctcgccgaccactaccagcagaacacccc
catcggcgacggccccgtgctgctgcccgacaaccactacctgagcacccagtccgccctgagcaaagaccccaacgagaagcgc
gatcacatggtcctgctggagttcgtgaccgccgccgggatcactctcggcatggacgagctgtacaagtaagtataccgataatcaa
cctctggattacaaaatttgtgaaagattgactggtattcttaactatgttgctccttttacgctatgtggatacgctgctttaatgcctttg
tatcatgctattgcttcccgtatggctttcattttctcctccttgtataaatcctggttgctgtctctttatgaggagttgtggcccgttgtca
ggcaacgtggcgtggtgtgcactgtgtttgctgacgcaacccccactggttggggcattgccaccacctgtcagctcctttccgggact
ttcgctttccccctccctattgccacggcggaactcatcgccgcctgccttgcccgctgctggacaggggctcggctgttgggcactga
caattccgtggtgttgtcggggaagctgacgtcctttccatggctgctcgcctgtgttgccacctggattctgcgcgggacgtccttctg
ctacgtcccttcggccctcaatccagcggaccttccttcccgcggcctgctgccggctctgcggcctcttccgcgtcttcgccttcgccc
tcagacgagtcggatctccctttgggccgcctccccgcatcggttaattaaacgggtggcatccctgtgacccctccccagtgcctctc
ctggccctggaagttgccactccagtgcccaccagccttgtcctaataaaattaagttgcatcattttgtctgactaggtgtccttctata
atattatggggtggaggggggtggtatggagcaaggggcaagttgggaagacaacctgtagggcctgcggggtctattgggaacca
agctggagtgcagtggcacaatcttggctcactgcaatctccgcctcctgggttcaagcgattctcctgcctcagcctcccgagttgttg
ggattccaggcatgcatgaccaggctcagctaatttttgtttttttggtagagacggggtttcaccatattggccaggctggtctccaact
cctaatctcaggtgatctacccaccttggcctcccaaattgctgggattacaggcgtgaaccactgctcccttccctgtccttatcgata
gatctaggaacccctagtgatggagttggccactccctctctgcgcgctcgctcgctcactgaggccgggcgaccaaaggtcgcccg
acgcccgggctttgcccgggcggcctcagtgagcgagcgagcgcgcagctgcctgcaggcagatcccctcgaggggtacccaact
ccatgctcaacagtccccaggtacagcccaccctgcgtcgcaaccaggaacagctctacagcttcctggagcgccactcgccctactt
ccgcagccacagtgcgcagattaggagcgccacttctttttgtcacttgaaaaacatgtaaaaataatgtactagagacactttcaataa
aggcaaatgcttttatttgtacactctcgggtgattatttacccccacccttgccgtctgcgccgtttaaaaatcaaaggggttctgccgc

gcatcgctatgcgccactggcagggacacgttgcgatactggtgtttagtgctccacttaaactcaggcacaaccatccgcggcagct
cggtgaagttttcactccacaggctgcgcaccatcaccaacgcgtttagcaggtcgggcgccgatatcttgaagtcgcagttggggcc
tccgccctgcgcgcgcgagttgcgatacacagggttgcagcactggaacactatcagcgccgggtggtgcacgctggccagcacgc
tcttgtcggagatcagatccgcgtccaggtcctccgcgttgctcagggcgaacggagtcaactttggtagctgccttcccaaaaaggg
cgcgtgcccaggctttgagttgcactcgcaccgtagtggcatcaaaaggtgaccgtgcccggtctgggcgttaggatacagcgcctg
cataaaagccttgatctgcttaaaagccacctgagcctttgcgccttcagagaagaacatgccgcaagacttgccggaaaactgattg
gccggacaggccgcgtcgtgcacgcagcaccttgcgtcggtgttggagatctgcaccacatttcggccccaccggttcttcacgatct
tggccttgctagactgctccttcagcgcgcgctgcccgttttcgctcgtcacatccatttcaatcacgtgctccttatttatcataatgctt
ccgtgtagacacttaagctcgccttcgatctcagcgcagcggtgcagccacaacgcgcagcccgtgggctcgtgatgcttgtaggtca
cctctgcaaacgactgcaggtacgcctgcaggaatcgccccatcatcgtcacaaaggtcttgttgctggtgaaggtcagctgcaaccc
gcggtgctcctcgttcagccaggtcttgcatacggccgccagagcttccacttggtcaggcagtagtttgaagttcgcctttagatcgtt
atccacgtggtacttgtccatcagcgcgcgcgcagcctccatgcccttctcccacgcagacacgatcggcacactcagcgggttcatc
accgtaatttcactttccgcttcgctgggctcttcctcttcctcttgcgtccgcataccacgcgccactgggtcgtcttcattcagccgcc
gcactgtgcgcttacctcctttgccatgcttgattagcaccggtgggttgctgaaacccaccatttgtagcgccacatcttctctttcttcc
tcgctgtccacgattacctctggtgatggcgggcgctcgggcttgggagaagggcgcttctttttcttcttgggcgcaatggccaaatc
cgccgccgaggtcgatggccgcgggctgggtgtgcgcggcaccagcgcgtcttgtgatgagtcttcctcgtcctcggactcgatacg
ccgcctcatccgctttttgggggcgcccggggaggcggcggcgacggggacggggacgacacgtcctccatggttgggggacgt
cgcgccgcaccgcgtccgcgctcgggggtggtttcgcgctgctcctcttcccgactggccatttccttctcctataggcagaaaaagat
catggagtcagtcgagaagaaggacagcctaaccgccccctctgagttcgccaccaccgcctccaccgatgccgccaacgcgccta
ccaccttccccgtcgaggcacccccgcttgaggaggaggaagtgattatcgagcaggacccaggttttgtaagcgaagacgacgag
gaccgctcagtaccaacagaggataaaaagcaagaccaggacaacgcagaggcaaacgaggaacaagtcgggcgggggggacga
aaggcatggcgactacctagatgtgggagacgacgtgctgttgaagcatctgcagcgccagtgcgccattatctgcgacgcgttgca
agagcgcagcgatgtgcccctcgccatagcggatgtcagccttgcctacgaacgccacctattctcaccgcgcgtacccccccaaacg
ccaagaaaacggcacatgcgagcccaacccgcgcctcaacttctaccccgtatttgccgtgccagaggtgcttgccacctatcacatc
tttttccaaaactgcaagatacccctatcctgccgtgccaaccgcagccgagcggacaagcagctggccttgcggcagggcgctgtc
atacctgatatcgcctcgctcaacgaagtgccaaaaatctttgagggtcttggacgcgacgagaagcgcgcggcaaacgctctgcaa
caggaaaacagcgaaaatgaaagtcactctggagtgttggtggaactcgagggtgacaacgcgcgcctagccgtactaaaacgcag
catcgaggtcacccactttgcctacccggcacttaacctacccccccaaggtcatgagcacagtcatgagtgagctgatcgtgcgccgt
gcgcagcccctggagagggatgcaaatttgcaagaacaaacagaggagggcctacccgcagttggcgacgagcagctagcgcgct
ggcttcaaacgcgcgagcctgccgacttggaggagcgacgcaaactaatgatggccgcagtgctcgttaccgtggagcttgagtgca
tgcagcggttctttgctgacccggagatgcagcgcaagctagaggaaacattgcactacacctttcgacagggctacgtacgccagg
cctgcaagatctccaacgtggagctctgcaacctggtctcctaccttggaattttgcacgaaaaccgccttgggcaaaacgtgcttcatt
ccacgctcaagggcgaggcgcgccgcgactacgtccgcgactgcgtttacttatttctatgctacacctggcagacggccatgggcgt
ttggcagcagtgcttggaggagtgcaacctcaaggagctgcagaaactgctaaagcaaaacttgaaggacctatggacggccttcaa
cgagcgctccgtggccgcgcacctggcggacatcattttccccgaacgcctgcttaaaaccctgcaacagggtctgccagacttcacc
agtcaaagcatgttgcagaactttaggaactttatcctagagcgctcaggaatcttgcccgccacctgctgtgcacttcctagcgacttt

gtgcccattaagtaccgcgaatgccctccgccgctttggggccactgctaccttctgcagctagccaactaccttgcctaccactctga
cataatggaagacgtgagcggtgacggtctactggagtgtcactgtcgctgcaacctatgcaccccgcaccgctccctggtttgcaat
tcgcagctgcttaacgaaagtcaaattatcggtacctttgagctgcagggtccctcgcctgacgaaaagtccgcggctccggggttga
aactcactccggggctgtggacgtcggcttaccttcgcaaatttgtacctgaggactaccacgcccacgagattaggttctacgaaga
ccaatcccgcccgcctaatgcggagcttaccgcctgcgtcattacccagggccacattcttggccaattgcaagccatcaacaaagcc
cgccaagagtttctgctacgaaagggacgggggtttacttggacccccagtccggcgaggagctcaacccaatccccccgccgcc
gcagccctatcagcagcagccgcgggcccttgcttcccaggatggcacccaaaaagaagctgcagctgccgccgccacccacggac
gaggaggaatactgggacagtcaggcagaggaggttttggacgaggaggaggaggacatgatggaagactgggagagcctagac
gaggaagcttccgaggtcgaagaggtgtcagacgaaacaccgtcaccctcggtcgcattcccctcgccggcgccccagaaatcggc
aaccggttccagcatggctacaacctccgctcctcaggcgccgccggcactgcccgttcgccgacccaaccgtagatgggacaccac
tggaaccagggccggtaagtccaagcagccgccgccgttagcccaagagcaacaacagcgccaaggctaccgctcatggcgcggg
cacaagaacgccatagttgcttgcttgcaagactgtgggggcaacatctccttcgcccgccgctttcttctctaccatcacggcgtggc
cttcccccgtaacatcctgcattactaccgtcatctctacagcccatactgcaccggcggcagcggcagcggcagcaacagcagcgg
ccacacagaagcaaaggcgaccggatagcaagactctgacaaagcccaagaaatccacagcggcggcagcagcaggaggagga
gcgctgcgtctggcgcccaacgaacccgtatcgacccgcgagcttagaaacaggatttttcccactctgtatgctatatttcaacagag
caggggccaagaacaagagctgaaaataaaaaacaggtctctgcgatccctcacccgcagctgcctgtatcacaaaagcgaagatc
agcttcggcgcacgctggaagacgcggaggctctcttcagtaaatactgcgcgctgactcttaaggactagtttcgcgccctttctcaa
atttaagcgcgaaaactacgtcatctccagcggccacacccggcgccagcacctgttgtcagcgccattatgagcaaggaaattccca
cgccctacatgtggagttaccagccacaaatgggacttgcggctggagctgcccaagactactcaacccgaataaactacatgagcg
cgggaccccacatgatatcccgggtcaacggaatacgcgcccaccgaaaccgaattctcctggaacaggcggctattaccaccacac
ctcgtaataaccttaatccccgtagttggcccgctgccctggtgtaccaggaaagtcccgctcccaccactgtggtacttcccagagac
gcccaggccgaagttcagatgactaactcaggggcgcagcttgcgggcggctttcgtcacagggtgcggtcgcccgggctgtacag
gcgcgccgaattcgttttagggcggagtaacttgtatgtgttgggaattgtagttttcttaaaatgggaagtgacgtaacgtgggaaaa
cggaagtgacgatttgaggaagttgtgggtttttggctttcgtttctgggcgtaggttcgcgtgcggttttctgggtgtttttgtggact
ttaaccgttacgtcatttttagtcctatatatactcgctctgcacttggccctttttacactgtgactgattgagctggtgccgtgtcgag
tggtgtttttttaataggttttctttttactggtaaggctgactgttatggctgccgctgtggaagcgctgtatgttgttctggagcggga
gggtgctattttgcctaggcaggagggtttttcaggtgtttatgtgtttttctctcctattaattttgttatacctcctatgggggctgtaatg
ttgtctctacgcctgcgggtatgtattcccccgggctatttcggtcgctttttagcactgaccgatgtgaatcaacctgatgtgtttaccg
agtcttacattatgactccggacatgaccgaggagctgtcggtggtgctttttaatcacggtgaccagtttttttacggtcacgccggca
tggccgtagtccgtcttatgcttataagggttgtttttcctgttgtaagacaggcttctaatgtttaaatgtttttttgttattttattttgtgtt
tatgcagaaacccgcagacatgtttgagagaaaaatggtgtctttttctgtggtggttccggagcttacctgcctttatctgcatgagcat
gactacgatgtgctttctttttgcgcgaggctttgcctgatttttgagcagcaccttgcattttatatcgccgcccatgcaacaagctta
catcggggctacgctggttagcatagctccgagtatgcgtgtcataatcagtgtgggttcttttgtcatggttcctggcggggaagtgg
ccgcgctggtccgtgcagacctgcacgattatgttcagctggccctgcgaagggacctacgggatcgcggtatttttgttaatgttccg
cttttgaatcttatacaggtctgtgaggaacctgaatttttgcaatcatgattcgctgcttgaggctgaaggtggagggcgctctggagc
agatttttacaatggccggacttaatattcgggatttgcttagagatatattgagaaggtggcgagatgagaattatttgggcatggttg

aaggtgctggaatgtttatagaggagattcaccctgaagggtttagcctttacgtccacttggacgtgagggccgtttgccttttggaa
gccattgtgcaacatcttacaaatgccattatctgttctttggctgtagagtttgaccacgccaccggaggggagcgcgttcacttaata
gatcttcattttgaggttttggataatcttttggaataaaaaaaaaaacatggttcttccagctcttcccgctcctcccgtgtgtgactcgc
agaacgaatgtgtaggttggctgggtgtggcttattctgcggtggtggatgttatcagggcagcggcgcatgaaggagtttacataga
acccgaagccagggggcgcctggatgctttgagagagtggatatactacaactactacacagagcgatctaagcggcgagaccgga
gacgcagatctgtttgtcacgcccgcacctggttttgcttcaggaaatatgactacgtccggcgttccatttggcatgacactacgacca
acacgatctcggttgtctcggcgcactccgtacagtagggatcgtctacctccttttgagacagaaacccgcgctaccatactggagg
atcatccgctgctgcccgaatgtaacactttgacaatgcacaacgtgagttacgtgcgaggtcttccctgcagtgtgggatttacgctg
attcaggaatgggttgttccctgggatatggttctaacgcgggaggagcttgtaatcctgaggaagtgtatgcacgtgtgcctgtgttg
tgccaacattgatatcatgacgagcatgatgatccatggttacgagtcctgggctctccactgtcattgttccagtcccggttccctgca
gtgtatagccggcgggcaggttttggccagctggtttaggatggtggtggatggcgccatgtttaatcagaggtttatatggtaccgg
gaggtggtgaattacaacatgccaaaagaggtaatgtttatgtccagcgtgtttatgaggggtcgccacttaatctacctgcgcttgtg
gtatgatggccacgtgggttctgtggtccccgccatgagctttggatacagcgccttgcactgtgggattttgaacaatattgtggtgct
gtgctgcagttactgtgctgatttaagtgagatcagggtgcgctgctgtgcccggaggacaaggcgccttatgctgcgggcggtgcg
aatcatcgctgaggagaccactgccatgttgtattcctgcaggacggagcggcggcggcagcagtttattcgcgcgctgctgcagca
ccaccgccctatcctgatgcacgattatgactctacccccatgtaggcgtggacttctccttcgccgcccgttaagcaaccgcaagttg
gacagcagcctgtggctcagcagctggacagcgacatgaacttaagtgagctgcccggggagtttattaatatcactgatgagcgttt
ggctcgacaggaaaccgtgtggaatataacacctaagaatatgtctgttacccatgatatgatgctttttaaggccagccggggagaa
aggactgtgtactctgtgtgttgggagggaggtggcaggttgaatactagggttctgtgagtttgattaaggtacggtgatctgtataa
gctatgtggtggtggggctatactactgaatgaaaaatgacttgaaattttctgcaattgaaaaataaacacgttgaaacataacacaa
acgattctttattcttgggcaatgtatgaaaaagtgtaagaggatgtggcaaatatttcattaatgtagttgtggccagaccagtcccat
gaaaatgacatagagtatgcacttggagttgtgtctcctgtttcctgtgtaccgtttagtaagcttccctcctgacgcggtaggaggagg
ggagggtgccctgcatgtctgccgctgctcttgctcttgccgctgctgaggagggggcgcatctgccgcagcaccggatgcatctg
ggaaaagcaaaaaggggctcgtccctgtttccggaggaatttgcaagcggggtcttgcatgacggggaggcaaacccccgttcgc
cgcagtccggccggtccgagactcgaaccgggggtcccgcgactcaacccttggaaaataaccctccggctacagggagcgagcc
acttaatgctttcgctttccagcctaaccgcttacgctgcgcgcggccagtggccaaaaaagctagcgcagcagccgccgcgcctgg
aaggaagccaaaaggagcactcccccgttgtctgacgtcgcacacctgggttcgacacgcgggcggtaaccgcatggatcacggcg
gacggccggatacggggctcgaaccccggtcgtccgccatgatacccttgcgaatttatccaccagaccacggaagagtgcccgctt
acaggctctccttttgcacggtctagagcgtcaacgattgcgcgcgcctgaccggccagagcgtcccgaccatggagcactttttgcc
gctgcgcaacatctggaaccgcgtccgcgactttccgcgcgcctccaccaccgccgccggcatcacctggatgtccaggtacatcta
ccaattgacatgtgagcaaaagggcgatcgctctagaactagtgtagccatggaaactagataagaaagaaatacgcagagaccaa
agttcaactgaaacgaattaaacggtttattgattaacaagcaattacaggggacgggtaaggtaacgggtgccaatggggcgaggc
tcagtataaagtccattgttgtccacagtaaaatcaacgttggcagattttgcataattggatgtgtactgcacttcgggattccagcgct
tgctgttttctttctgcagctcccattcaatttccacactcacttgtcctgtggagtattgggtgatgaatgaagcaaactttgtagctgaa
aactccgccggaggattcgcaggaacaggcgtgtttttgatgaggatctgaggaggcgggttcttgagtccaaagccgcccataaga
ggagacgggtgaaagtgtccatctgtgtgaggaattttggcccaaatgggaccctgcaggtacacgtctctatcttgccacaccatgc

caggtaatgctcccatagcatgcacatctccggtcgcagggtctgtgctgctgctctggaaattgactgccacggtcccaaatctttcg
gtggccacagggttagtggctttaatttcctcttcgtctgtaatcatgacattgtccaatgcagtgtttgaagctccggcgctctcttttcc
aaaaatcatgacaccgctcatgggaaagaacttgtcttcgtcgtctttgtgtgaggccatagcagtgccagggttgatgatggattcac
gcccattgaggttatattttgaagcaccagtccaggtaaaattgctgttgttgttgtctgtttttgttttagaaacgcgctgctgccgataa
cagggtccaggtagccagtttttgggctgaacagacatgccagctggagacccacggctaaacagcaagtccttgttttgggcacttc
cggactgattttgagttctgttcaggtaatacaggtattggtcgatgagaggattcatcagccggtccaggctctggctgtgcgcgtag
ctgctgtggaaaggcacttcctcaaaggtgtagctgaaggtaaagttgttgcccgttctcagcatctgagaagggaaatattccaggc
agtaaaaggatgaacgtcccacggcttggctgccattgttgagcgtcaggtagccgtattgcggaatcatgaacacgtccgccggga
acggagggaggcagccctggtgcgcagagccgaggacgtacggaagctggtactccgagtccgagaagacttgaaccgtgctggt
aaggttattagcgatggttgtgacgccatcattcgtcgtgacctccttgacttggatgttgaagagtttgaagttgagtctcttgggccg
gaatccccaattgttgttgatgagtcgctgccagtcacgtggtgaaaagtggcagtggaatctgttgaaatcaaaatacccccagggg
gtgctgtagccgaagtagtggttgtcgttgctggcccccgttgaagcactggagatttgcttgtagaggtggttattgtaggtgggcaa
ggcccaggtgcgggtgctggtggtgatgactctgtcgcccagccatgtggaatcgcaatgccaatttcctgaggcattacccactccg
tcggcgccttcgttattgtctgccattggtgcgccaccgcctgaagccattgtagtaggtcccacagcagcgggggttgctggaggttc
tccgagaggttgtggatcggggactgactctgagtcgccagtctgaccaaaattgagtctctttttagcgggctgctggcctgtcttgc
cgatgcccgaggaggagtctggctcttgtggcgactgctctaccggacgtttctttccaggagccgtcttagcgccttcctcaaccaga
ccgagaggttcgagaacccgcttcttggcctggaagactgctcgcccgaggttgcccccaaaagacgtatcttcttgcagacgctcct
gaaactcggcgtcggcgtggttataccgcaggtacggattgtcacccgctttgagctgctggtcgtaggccttgtcgtgctcgagggc
cgctgcgtccgccgcgttgacgggctccccttgtcgagtccgttgaagggtccgaggtacttgtagccaggaagcaccagaccccg
gccgtcgtcctgcttttgctggttggctttgggcttcggggctccaggtttcaagtcccaccactcgcgaatgccctcagagaggttgtc
ctcgagccaatctggaagataaccatcggcagccatacctgatttaaatcatttattgttcaaagatgcagtcatccaaatccacattga
ccagatcgcaggcagtgcaagcgtctggcacctttcccatgatatgatgaatgtagcacagtttctgatacgcctttttgacgacagaa
acgggttgagattctgacacgggaaagcactctaaacagtctttctgtccgtgagtgaagcagatatttgaattctgattcattctctcg
cattgtctgcagggaaacagcatcagattcatgcccacgtgacgagaacatttgttttggtacctgtctgcgtagttgatcgaagcttcc
gcgtctgacgtcgatggctgcgcaactgactcgcgcacccgtttgggctcacttatatctgcgtcactgggggcgggtcttttcttggc
tccacccttttgacgtagaattcatgctccacctcaaccacgtgatcctttgcccaccggaaaaagtctttgacttcctgcttggtgacc
ttcccaaagtcatgatccagacggcgggtgagttcaaatttgaacatccggtcttgcaacggctgctggtgttcgaaggtcgttgagtt
cccgtcaatcacggcgcacatgttggtgttggaggtgacgatcacgggagtcgggtctatctgggccgaggacttgcatttctggtcc
acgcgcaccttgcttcctccgagaatggctttggccgactccacgaccttggcggtcatcttcccctcctcccaccagatcaccatcttg
tcgacacagtcgttgaagggaaagttctcattggtccagtttacgcacccgtagaagggcacagtgtgggctatggcctccgcgatgt
tggtcttcccggtagttgcaggcccaaacagccagatggtgttcctcttgccgaactttttcgtggcccatcccagaaagacggaagcc
gcatattggggatcgtacccgtttagttccaaaattttataaatccgattgctggaaatgtcctccacgggctgctggcccaccaggtag
tcgggggcggttttagtcaggctcataatctttcccgcattgtccaaggcagccttgatttgggaccgcgagttggaggccgcattgaa
ggagatgtatgaggcctggtcctcctggatccactgcttctccgaggtaatccccttgtccacgagccacccgaccagctccatgtacc
tggctgaagtttttgatctgatcaccggcgcatcagaattgggattctgattctctttgttctgctcctgcgtctgcgacacgtgcgtcag
atgctgcgccaccaaccgtttacgctccgtgagattcaaacaggcgcttaaatactgttccatattagtccacgcccactggagctcag

gctgggttttggggagcaagtaattggggatgtagcactcatccaccaccttgttcccgcctccggcgccatttctggtctttgtgaccg
cgaaccagtttggcaaagtcggctcgatcccgcggtaaattctctgaatcagtttttcgcgaatctgactcaggaaacgtcccaaaacc
atggatttcaccccggtggtttccacgagcacgtgcatgtggaagtagctctctcccttctcaaattgcacaaagaaaagggcctccgg
ggccttactcacacggcgccattccgtcagaaagtcgcgctgcagcttctcggccacggtcaggggtgcctgctcaatcagattcaga
tccatgtcagaatctggcggcaactcccattccttctcggccacccagttcacaaagctgtcagaaatgccgggcagatgctcgtcaa
ggtcgctggggaccttaatcacaatctcgtaaaaccccggcatggcggctgcgcgttcaaacctcccgcttcaaaatggagaccctgc
gtgctcactcgggcttaaatacccagcgtgaccacatggtgtcgcaaaatgtcgcaaaacactcacgtgacctctaatacaggacctc
cctaaccctatgacgtaattcacgtcacgactccacccctccgcggccgcgtttaaactagtgtggcggaagtgtgatgttgcaagtgt
ggcggaacacatgtaagcgacggatgtggcaaaagtgacgtttttggtgtgcgccggtgtacacaggaagtgacaattttcgcgcgg
ttttaggcggatgttgtagtaaatttgggcgtaaccgagtaagatttggccattttcgcgggaaaactgaataagaggaagtgaaatct
gaataattttgtgttactcatagcgcgtaatatttgtctagggccgcggggactttgaccgtttacgtggagactcgcccaggtgtttttc
tcaggtgttttccgcgttccgggtcaaagttggcgttttattattatagtcagctgacgtgtagtgtatttatacccggtgagttcctcaag
aggccactcttgagtgccagcgagtagagttttctcctccgagccgctccgacaccgggactgaaaatgagacatattatctgccacg
gaggtgttattaccgaagaaatggccgccagtcttttggaccagctgatcgaagaggtactggctgataatcttccacctcctagccat
tttgaaccacctacccttcacgaactgtatgatttagacgtgacggcccccgaagatcccaacgaggaggcggtttcgcagatttttcc
cgactctgtaatgttggcggtgcaggaagggattgacttactcacttttccgccggcgcccggttctccggagccgcctcacctttccc
ggcagcccgagcagccggagcagagagccttgggtccggtttctatgccaaaccttgtaccggaggtgatcgatcttacctgccacg
aggctggctttccacccagtgacgacgaggatgaagagggtgaggagtttgtgttagattatgtggagcaccccgggcacggttgca
ggtcttgtcattatcaccggaggaatacgggggacccagatattatgtgttcgctttgctatatgaggacctgtggcatgtttgtctaca
gtaagtgaaaattatgggcagtgggtgatagagtggtgggtttggtgtggtaattttttttttaatttttacagttttgtggtttaaagaatt
ttgtattgtgatttttttaaaaggtcctgtgtctgaacctgagcctgagcccgagccagaaccggagcctgcaagacctacccgccgtc
ctaaaatggcgcctgctatcctgagacgcccgacatcacctgtgtctagagaatgcaatagtagtacggatagctgtgactccggtcct
tctaacacacctcctgagatacacccggtggtcccgctgtgccccattaaaccagttgccgtgagagttggtgggcgtcgccaggctg
tggaatgtatcgaggacttgcttaacgagcctgggcaacctttggacttgagctgtaaacgccccaggccataaggtgtaaacctgtg
attgcgtgtgtggttaacgcctttgtttgctgaatgagttgatgtaagtttaataaagggtgagataatgtttaacttgcatggcgtgttaa
atggggcggggcttaaagggtatataatgcgccgtgggctaatcttggttacatctgacctcatggaggcttgggagtgtttggaaga
tttttctgctgtgcgtaacttgctggaacagagctctaacagtacctcttggttttggaggtttctgtggggctcatcccaggcaaagtta
gtctgcagaattaaggaggattacaagtgggaatttgaagagcttttgaaatcctgtggtgagctgtttgattctttgaatctgggtcac
caggcgcttttccaagagaaggtcatcaagactttggatttttccacaccggggcgcgctgcggctgctgttgctttttgagttttataa
aggataaatggagcgaagaaacccatctgagcggggggtacctgctggattttctggccatgcatctgtggagagcggttgtgagac
acaagaatcgcctgctactgttgtcttccgtccgcccggcgataataccgacggaggagcagcagcagcaggaggaagccagg
cggcggcggcaggagcagagcccatggaacccgagagccggcctggaccctcgggaatgaatgttgtacaggtggctgaactgta
tccagaactgagacgcattttgacaattacagaggatgggcaggggctaaagggggtaaagagggagcgggggcttgtgaggct
acagaggaggctaggaatctagcttttagcttaatgaccagacaccgtcctgagtgtattacttttcaacagatcaaggataattgcgct
aatgagcttgatctgctggcgcagaagtattccatagagcagctgaccacttactggctgcagccaggggatgattttgaggaggcta
ttagggtatatgcaaaggtggcacttaggccagattgcaagtacaagatcagcaaacttgtaaatatcaggaattgttgctacatttctg

ggaacggggccgaggtggagatagatacggaggatagggtggcctttagatgtagcatgataaatatgtggccgggggtgcttggc
atggacggggtggttattatgaatgtaaggtttactggccccaattttagcggtacggttttcctggccaataccaaccttatcctacac
ggtgtaagcttctatgggtttaacaatacctgtgtggaagcctggaccgatgtaagggttcggggctgtgccttttactgctgctggaa
gggggtggtgtgtcgccccaaaagcagggcttcaattaagaaatgcctctttgaaaggtgtaccttgggtatcctgtctgagggtaact
ccagggtgcgccacaatgtggcctccgactgtggttgcttcatgctagtgaaaagcgtggctgtgattaagcataacatggtatgtgg
caactgcgaggacagggcctctcagatgctgacctgctcggacggcaactgtcacctgctgaagaccattcacgtagccagccactc
tcgcaaggcctggccagtgtttgagcataacatactgacccgctgttccttgcatttgggtaacaggagggggtgttcctaccttacc
aatgcaatttgagtcacactaagatattgcttgagcccgagagcatgtccaaggtgaacctgaacggggtgtttgacatgaccatgaa
gatctggaaggtgctgaggtacgatgagacccgcaccaggtgcagaccctgcgagtgtggcggtaaacatattaggaaccagcctg
tgatgctggatgtgaccgaggagctgaggcccgatcacttggtgctggcctgcacccgcgctgagtttggctctagcgatgaagatac
agattgaggtactgaaatgtgtgggcgtggcttaagggtgggaaagaatatataaggtgggggtcttatgtagttttgtatctgttttgc
agcagccgccgccgccatgagcaccaactcgtttgatggaagcattgtgagctcatatttgacaacgcgcatgcccccatgggccgg
ggtgcgtcagaatgtgatgggctccagcattgatggtcgccccgtcctgcccgcaaactctactaccttgacctacgagaccgtgtctg
gaacgccgttggagactgcagcctccgccgccgcttcagccgctgcagccaccgcccgcgggattgtgactgactttgctttcctgag
cccgcttgcaagcagtgcagcttcccgttcatccgcccgcgatgacaagttgacggctcttttggcacaattggattctttgacccggg
aacttaatgtcgtttctcagcagctgttggatctgcgccagcaggtttctgccctgaaggcttcctcccctcccaatgcggtttaaaacat
aaataaaaaaccagactctgtttggatttggatcaagcaagtgtcttgctgtctttatttaggggttttgcgcgcgcggtaggcccggga
ccagcggccggccgactcgctgcgctcggtcgttcgggtaaagcctggggtgcctaatgagcaaaaggccagcaaaaggccagga
accgtaaaaaggccgcgttgctggcgtttttccataggctccgcccccctgacgagcatcacaaaaatcgacgctcaagtcagaggtg
gcgaaacccgacaggactataaagataccaggcgtttccccctggaagctccctcgtgcgctctcctgttccgaccctgccgcttacc
ggatacctgtccgcctttctcccttcgggaagcgtggcgctttctcatagctcacgctgtaggtatctcagttcggtgtaggtcgttcgct
ccaagctgggctgtgtgcacgaaccccccgttcagcccgaccgctgcgccttatccggtaactatcgtcttgagtccaacccggtaag
acacgacttatcgccactggcagcagccactggtaacaggattagcagagcgaggtatgtaggcggtgctacagagttcttgaagtg
gtggcctaactacggctacactagaagaacagtatttggtatctgcgctctgctgaagccagttaccttcggaaaaagagttggtagct
cttgatccggcaaacaaaccaccgctggtagcggtggtttttttgtttgcaagcagcagattacgcgcagaaaaaaaggatctcaaga
agatcctttgatcttttctacggggtctgacgctcagtggaacgaaaactcacgttaagggattttggtcatgagattatcaaaaaggat
cttcacctagatccttttaaattaaaaatgaagttttaaatcaatctaaagtatatatgagtaaacttggtctgacagttattagaaaaattc
atccagcagacgataaaacgcaatacgctggctatccggtgccgcaatgccatacagcaccagaaaacgatccgcccattcgccgcc
cagttcttccgcaatatcacgggtggccagcgcaatatcctgataacgatccgccacgcccagacggccgcaatcaataaagccgct
aaaacggccattttccaccataatgttcggcaggcacgcatcaccatgggtcaccaccagatcttcgccatccggcatgctcgctttca
gacgcgcaaacagctctgccggtgccaggccctgatgttcttcatccagatcatcctgatccaccaggcccgcttccatacgggtacg
cgcacgttcaatacgatgtttcgcctgatgatcaaacggacaggtcgccgggtccagggtatgcagacgacgcatggcatccgccat
aatgctcactttttctgccggcgccagatggctagacagcagatcctgacccggcacttcgcccagcagcagccaatcacggcccgct
tcggtcaccacatccagcaccgccgcacacggaacaccggtggtggccagccagctcagacgcgccgcttcatcctgcagctcgttc
agcgcaccgctcagatcggttttcacaaacagcaccggacgaccctgcgcgctcagacgaaacaccgccgcatcagagcagccaat
ggtctgctgcgcccaatcatagccaaacagacgttccacccacgctgccgggctacccgcatgcaggccatcctgttcaatcatactc

ttcctttttcaatattattgaagcatttatcagggttattgtctcatgagcggatacatatttgaatgtatttagaaaaataaacaaataggg
gttccgcgcacatttccccgaaaagtgccacctaaattgtaagcgttaatattttgttaaaattcgcgttaaatttttgttaaatcagctcat
tttttaaccaataggtttattctgtcttttatttcaggcaccgggcttgcgggtcatgcaccaggtgcgcggtccttcgggcacctcgac
gtcggcggtgacggtgaagccgagccgctcgtagaaggggaggttgcggggcgcggaggtctccaggaaggcgggcaccccggc
gcgctcggccgcctccactccggggagcacgacggcgctgcccagacccttgccctggtggtcgggcgagacgccgacggtggcc
aggaaccacgcgggctccttgggccggtgcggcgccaggaggccttccatctgttgctgcgcggccagccgggaaccgctcaactc
ggccatgcgcgggccgatctcggcgaacaccgcccccgcttcgacgctctccggcgtggtccagaccgccaccgcggcgccgtcgt
ccgcgacccacaccttgccgatgtcgagcccgacgcgcgtgaggaagagttcttgcagctcggtgacccgctcgatgtggcggtcc
ggatcgacggtgtggcgcgtggcggggtagtcggcgaacgcggcggcgagggtgcgtacggccctggggacgtcgtcgcgggtg
gcgaggcgcaccgtgggcttgtactcggtcatggtggccacgtgaggtcgaaaggcccggagatgaggaagaggagaacagcgc
ggcagacgtgcgcttttgaagcgtgcagaatgccgggcctccggaggaccttcgggcgcccgccccgcccctgagcccgcccctga
gcccgcccccggacccaccccttcccagcctctgagcccagaaagcgaaggagcaaagctgctattggccgctgccccaaaggcct
acccgcttccattgctcagcggtgctgtccatctgcacgagactagtgagacgtgctacttccatttgtcacgtcctgcacgacgcgag
ctgcggggcggggggggaacttcctgactaggggaggagtagaaggtggcgcgaaggggccaccaaagaacggagccggttggc
gcctaccggtggatgtggaatgtgtgcgaggccagaggccacttgtgtagcgccaagtgcccagcggggctgctaaagcgcatgct
ccagactgccttgggaaaagcgcctcccctaccctcaagaattctcatgtttgacagcttatcatcgataagcttg

Supplemental data 5

Sequence of SYNp1088

ggtacccaactccatgctcaacagtccccaggtacagcccaccctgcgtcgcaaccaggaacagctctacagcttcctggagcgcca
ctcgccctacttccgcagccacagtgcgcagattaggagcgccacttctttttgtcacttgaaaaacatgtaaaaataatgtactagaga
cactttcaataaaggcaaatgcttttatttgtacactctcgggtgattatttacccccacccttgccgtctgcgccgtttaaaaatcaaag
gggttctgccgcgcatcgctatgcgccactggcagggacacgttgcgatactggtgtttagtgctccacttaaactcaggcacaacca
tccgcggcagctcggtgaagttttcactccacaggctgcgcaccatcaccaacgcgtttagcaggtcgggcgccgatatcttgaagtc
gcagttggggcctccgccctgcgcgcgcgagttgcgatacacagggttgcagcactggaacactatcagcgccgggtggtgcacgc
tggccagcacgctcttgtcggagatcagatccgcgtccaggtcctccgcgttgctcagggcgaacggagtcaactttggtagctgcct
tcccaaaaagggcgcgtgcccaggctttgagttgcactcgcaccgtagtggcatcaaaaggtgaccgtgcccggtctgggcgttagg
atacagcgcctgcataaaagccttgatctgcttaaaagccacctgagcctttgcgccttcagagaagaacatgccgcaagacttgccg
gaaaactgattggccggacaggccgcgtcgtgcacgcagcaccttgcgtcggtgttggagatctgcaccacatttcggccccaccgg
ttcttcacgatcttggccttgctagactgctccttcagcgcgcgctgcccgttttcgctcgtcacatccatttcaatcacgtgctccttattt
atcataatgcttccgtgtagacacttaagctcgccttcgatctcagcgcagcggtgcagccacaacgcgcagcccgtgggctcgtgat
gcttgtaggtcacctctgcaaacgactgcaggtacgcctgcaggaatcgccccatcatcgtcacaaaggtcttgttgctggtgaaggt
cagctgcaacccgcggtgctcctcgttcagccaggtcttgcatacggccgccagagcttccacttggtcaggcagtagtttgaagttc
gcctttagatcgttatccacgtggtacttgtccatcagcgcgcgcgcagcctccatgcccttctcccacgcagacacgatcggcacact
cagcgggttcatcaccgtaatttcactttccgcttcgctgggctcttcctcttcctcttgcgtccgcataccacgcgccactgggtcgtct
tcattcagccgccgcactgtgcgcttacctcctttgccatgcttgattagcaccggtgggttgctgaaacccaccatttgtagcgccaca
tcttctctttcttcctcgctgtccacgattacctctggtgatggcgggcgctcgggcttgggagaagggcgcttctttttcttcttgggcg
caatggccaaatccgccgccgaggtcgatggccgcgggctgggtgtgcgcggcaccagcgcgtcttgtgatgagtcttcctcgtcct
cggactcgatacgccgcctcatccgcttttttgggggcgcccggggaggcggcggcgacggggacggggacgacacgtcctccat
ggttgggggacgtcgcgccgcaccgcgtccgcgctcgggggtggtttcgcgctgctcctcttcccgactggccatttccttctcctata
ggcagaaaaagatcatggagtcagtcgagaagaaggacagcctaaccgcccccctctgagttcgccaccaccgcctccaccgatgcc
gccaacgcgcctaccaccttccccgtcgaggcaccccccgcttgaggaggaggaagtgattatcgagcaggacccaggttttgtaagc
gaagacgacgaggaccgctcagtaccaacagaggataaaaagcaagaccaggacaacgcagaggcaaacgaggaacaagtcgg
gcgggggacgaaaggcatggcgactacctagatgtgggagacgacgtgctgttgaagcatctgcagcgccagtgcgccattatct
gcgacgcgttgcaagagcgcagcgatgtgcccctcgccatagcggatgtcagccttgcctacgaacgccacctattctcaccgcgcg
taccccccaaacgccaagaaaacggcacatgcgagcccaacccgcgcctcaacttctaccccgtatttgccgtgccagaggtgcttg
ccacctatcacatctttttccaaaactgcaagatacccctatcctgccgtgccaaccgcagccgagcggacaagcagctggccttgcg
gcagggcgctgtcatacctgatatcgcctcgctcaacgaagtgccaaaaatctttgagggtcttggacgcgacgagaagcgcgcggc
aaacgctctgcaacaggaaaacagcgaaaatgaaagtcactctggagtgttggtggaactcgagggtgacaacgcgcgcctagccg
tactaaaacgcagcatcgaggtcacccactttgcctacccggcacttaacctacccccccaaggtcatgagcacagtcatgagtgagct
gatcgtgcgccgtgcgcagcccctggagagggatgcaaatttgcaagaacaaacagaggagggcctacccgcagttggcgacgag
cagctagcgcgctggcttcaaacgcgcgagcctgccgacttggaggagcgacgcaaactaatgatggccgcagtgctcgttaccgt
ggagcttgagtgcatgcagcggttctttgctgacccggagatgcagcgcaagctagaggaaacattgcactacacctttcgacaggg

ctacgtacgccaggcctgcaagatctccaacgtggagctctgcaacctggtctcctaccttggaattttgcacgaaaaccgccttgggc
aaaacgtgcttcattccacgctcaagggcgaggcgcgccgcgactacgtccgcgactgcgtttacttatttctatgctacacctggcag
acggccatgggcgtttggcagcagtgcttggaggagtgcaacctcaaggagctgcagaaactgctaaagcaaaacttgaaggacct
atggacggccttcaacgagcgctccgtggccgcgcacctggcggacatcattttccccgaacgcctgcttaaaaccctgcaacaggg
tctgccagacttcaccagtcaaagcatgttgcagaactttaggaactttatcctagagcgctcaggaatcttgcccgccacctgctgtgc
acttcctagcgactttgtgcccattaagtaccgcgaatgccctccgccgctttggggccactgctaccttctgcagctagccaactacct
tgcctaccactctgacataatggaagacgtgagcggtgacggtctactggagtgtcactgtcgctgcaacctatgcaccccgcaccgc
tccctggtttgcaattcgcagctgcttaacgaaagtcaaattatcggtacctttgagctgcagggtccctcgcctgacgaaaagtccgc
ggctccggggttgaaactcactccggggctgtggacgtcggcttaccttcgcaaatttgtacctgaggactaccacgcccacgagatt
aggttctacgaagaccaatcccgcccgccaaatgcggagcttaccgcctgcgtcattacccagggccacattcttggccaattgcaag
ccatcaacaaagcccgccaagagtttctgctacgaaagggacgggggtttacttggacccccagtccggcgaggagctcaaccca
atccccccgccgccgcagccctatcagcagcagccgcgggcccttgcttcccaggatggcacccaaaaagaagctgcagctgccgc
cgccacccacggacgaggaggaatactgggacagtcaggcagaggaggttttggacgaggaggaggaggacatgatggaagact
gggagagcctagacgaggaagcttccgaggtcgaagaggtgtcagacgaaacaccgtcaccctcggtcgcattcccctcgccggcg
ccccagaaatcggcaaccggttccagcatggctacaacctccgctcctcaggcgccgccggcactgcccgttcgccgacccaaccgt
agatgggacaccactggaaccagggccggtaagtccaagcagccgccgccgttagcccaagagcaacaacagcgccaaggctacc
gctcatggcgcgggcacaagaacgccatagttgcttgcttgcaagactgtgggggcaacatctccttcgcccgccgctttcttctctac
catcacggcgtggccttcccccgtaacatcctgcattactaccgtcatctctacagcccatactgcaccggcggcagcggcagcggca
gcaacagcagcggccacacagaagcaaaggcgaccggatagcaagactctgacaaagcccaagaaatccacagcggcggcagca
gcaggaggaggagcgctgcgtctggcgcccaacgaacccgtatcgacccgcgagcttagaaacaggatttttcccactctgtatgct
atatttcaacagagcaggggccaagaacaagagctgaaaataaaaaacaggtctctgcgatccctcacccgcagctgcctgtatcac
aaaagcgaagatcagcttcggcgcacgctggaagacgcggaggctctcttcagtaaatactgcgcgctgactcttaaggactagtttc
gcgccctttctcaaatttaagcgcgaaaactacgtcatctccagcggccacacccggcgccagcacctgtcgtcagcgccattatgag
caaggaaattcccacgccctacatgtggagttaccagccacaaatgggacttgcggctggagctgcccaagactactcaacccgaat
aaactacatgagcgcgggaccccacatgatatcccgggtcaacggaatccgcgcccaccgaaaccgaattctcttggaacaggcgg
ctattaccaccacacctcgtaataaccttaatccccgtagttggcccgctgccctggtgtaccaggaaagtcccgctcccaccactgtg
gtacttcccagagacgcccaggccgaagttcagatgactaactcaggggcgcagcttgcgggcggctttcgtcacagggtgcggtcg
cccgggcgttttagggcggagtaacttgtatgtgttgggaattgtagttttcttaaaatgggaagttacgtaacgtgggaaaacggaag
tgacgatttgaggaagttgtgggtttttggctttcgtttctgggcgtaggttcgcgtgcggttttctgggtgtttttgtggactttaaccg
ttacgtcatttttagtcctatatatactcgctctgcacttggccctttttacactgtgactgattgagctggtgccgtgtcgagtggtgttt
ttttaataggttttctttttactggtaaggctgactgttatggctgccgctgtggaagcgctgtatgttgttctggagcgggagggtgct
attttgcctaggcaggagggtttttcaggtgtttatgtgtttttctctcctattaattttgttatacctcctatgggggctgtaatgttgtctct
acgcctgcgggtatgtattcccccgggctatttcggtcgctttttagcactgaccgatgtgaatcaacctgatgtgtttaccgagtcttac
attatgactccggacatgaccgaggagctgtcggtggtgctttttaatcacggtgaccagtttttttacggtcacgccggcatggccgta
gtccgtcttatgcttataagggttgtttttcctgttgtaagacaggcttctaatgtttaaatgtttttttgttattttattttgtgtttatgcaga
aacccgcagacatgtttgagagaaaaatggtgtctttttctgtggtggttccggagcttacctgcctttatctgcatgagcatgactacg

atgtgctttctttttgcgcgaggctttgcctgatttttgagcagcaccttgcattttatatcgccgcccatgcaacaagcttacatcggg
gctacgctggttagcatagctccgagtatgcgtgtcataatcagtgtgggttcttttgtcatggttcctggcggggaagtggccgcgct
ggtccgtgcagacctgcacgattatgttcagctggccctgcgaagggacctacgggatcgcggtatttttgttaatgttccgcttttgaa
tcttatacaggtctgtgaggaacctgaatttttgcaatcatgattcgctgcttgaggctgaaggtggagggcgctctggagcagattttt
acaatggccggacttaatattcgggatttgcttagagatatattgagaaggtggcgagatgagaattatttgggcatggttgaaggtgc
tggaatgtttatagaggagattcaccctgaagggtttagcctttacgtccacttggacgtgagggccgtttgccttttggaagccattgt
gcaacatcttacaaatgccattatctgttctttggctgtagagtttgaccacgccaccggaggggagcgcgttcacttaatagatcttca
ttttgaggttttggataatcttttggaataaaaaaaaaaacatggttcttccagctcttcccgctcctcccgtgtgtgactcgcagaacga
atgtgtaggttggctgggtgtggcttattctgcggtggtggatgttatcagggcagcggcgcatgaaggagtttacatagaacccgaa
gccagggggcgcctggatgctttgagagagtggatatactacaactactacacagagcgatctaagcggcgagaccggagacgcag
atctgtttgtcacgcccgcacctggttttgcttcaggaaatatgactacgtccggcgttccatttggcatgacactacgaccaacacgat
ctcggttgtctcggcgcactccgtacagtagggatcgtctacctccttttgagacagaaacccgcgctaccatactggaggatcatccg
ctgctgcccgaatgtaacactttgacaatgcacaacgtgagttacgtgcgaggtcttccctgcagtgtgggatttacgctgattcagga
atgggttgttccctgggatatggttctaacgcgggaggagcttgtaatcctgaggaagtgtatgcacgtgtgcctgtgttgtgccaaca
ttgatatcatgacgagcatgatgatccatggttacgagtcctgggctctccactgtcattgttccagtcccggttccctgcagtgtatagc
cggcgggcaggttttggccagctggtttaggatggtggtggatggcgccatgtttaatcagaggtttatatggtaccgggaggtggtg
aattacaacatgccaaaagaggtaatgtttatgtccagcgtgtttatgaggggtcgccacttaatctacctgcgcttgtggtatgatggc
cacgtgggttctgtggtccccgccatgagctttggatacagcgccttgcactgtgggattttgaacaatattgtggtgctgtgctgcagt
tactgtgctgatttaagtgagatcagggtgcgctgctgtgcccggaggacaaggcgccttatgctgcgggcggtgcgaatcatcgctg
aggagaccactgccatgttgtattcctgcaggacggagcggcggcggcagcagtttattcgcgcgctgctgcagcaccaccgcccta
tcctgatgcacgattatgactctacccccatgtaggcgtggacttctccttcgccgcccgttaagcaaccgcaagttggacagcagcct
gtggctcagcagctggacagcgacatgaacttaagtgagctgcccggggagtttattaatatcactgatgagcgtttggctcgacagg
aaaccgtgtggaatataacacctaagaatatgtctgttacccatgatatgatgctttttaaggccagccggggagaaaggactgtgtac
tctgtgtgttgggagggaggtggcaggttgaatactagggttctgtgagtttgattaaggtacggtgatctgtataagctatgtggtggt
ggggctatactactgaatgaaaaatgacttgaaattttctgcaattgaaaaataaacacgttgaaacataacacaaacgattctttattct
tgggcaatgtatgaaaaagtgtaagaggatgtggcaaatatttcattaatgtagttgtggccagaccagtcccatgaaaatgacataga
gtatgcacttggagttgtgtctcctgtttcctgtgtaccgtttagtccctcctgacgcggtaggaggaggggagggtgccctgcatgtct
gccgctgctcttgctcttgccgctgctgaggagggggcgcatctgccgcagcaccggatgcatctgggaaaagcaaaaaaggggc
tcgtccctgtttccggaggaatttgcaagcggggtcttgcatgacggggaggcaaacccccgttcgccgcagtccggccggtccgag
actcgaaccgggggtcccgcgactcaacccttggaaaataaccctccggctacagggagcgagccacttaatgctttcgctttccagc
ctaaccgcttacgctgcgcgcggccagtggccaaaaaagctagcgcagcagccgccgcgcctggaaggaagccaaaaggagcact
cccccgttgtctgacgtcgcacacctgggttcgacacgcgggcggtaaccgcatggatcacggcggacggccggatacggggctcg
aaccccggtcgtccgccatgatacccttgcgaatttatccaccagaccacggaagagtgcccgcttacaggctctccttttgcacggtc
tagagcgtcaacgattgcgcgcgcctgaccggccagagcgtcccgaccatggagcactttttgccgctgcgcaacatctggaaccgc
gtccgcgactttccgcgcgcctccaccaccgccgccggcatcacctggatgtccaggtacatctacctaatgagcaaaaggccagca
aaaggccaggaaccgtaaaaaggccgcgttgctggcgtttttccataggctccgcccccctgacgagcatcacaaaaatcgacgctc

aagtcagaggtggcgaaacccgacaggactataaagataccaggcgtttccccctggaagctccctcgtgcgctctcctgttccgacc
ctgccgcttaccggatacctgtccgcctttctcccttcgggaagcgtggcgctttctcatagctcacgctgtaggtatctcagttcggtgt
aggtcgttcgctccaagctgggctgtgtgcacgaaccccccgttcagcccgaccgctgcgccttatccggtaactatcgtcttgagtcc
aacccggtaagacacgacttatcgccactggcagcagccactggtaacaggattagcagagcgaggtatgtaggcggtgctacaga
gttcttgaagtggtggcctaactacggctacactagaagaacagtatttggtatctgcgctctgctgaagccagttaccttcggaaaaa
gagttggtagctcttgatccggcaaacaaaccaccgctggtagcggtggtttttttgtttgcaagcagcagattacgcgcagaaaaaa
aggatctcaagaagatcctttgatcttttctacggggtctgacgctcagtggaacgaaaactcacgttaagggattttggtcatgagatt
atcaaaaaggatcttcacctagatccttttaaattaaaaatgaagttttaaatcaatctaaagtatatatgagtaaacttggtctgacagtt
attagaaaaattcatccagcagacgataaaacgcaatacgctggctatccggtgccgcaatgccatacagcaccagaaaacgatccg
cccattcgccgcccagttcttccgcaatatcacgggtggccagcgcaatatcctgataacgatccgccacgcccagacggccgcaatc
aataaagccgctaaaacggccattttccaccataatgttcggcaggcacgcatcaccatgggtcaccaccagatcttcgccatccggc
atgctcgctttcagacgcgcaaacagctctgccggtgccaggccctgatgttcttcatccagatcatcctgatccaccaggcccgcttc
catacgggtacgcgcacgttcaatacgatgtttcgcctgatgatcaaacggacaggtcgccgggtccagggtatgcagacgacgcat
ggcatccgccataatgctcactttttctgccggcgccagatggctagacagcagatcctgacccggcacttcgcccagcagcagccaa
tcacggcccgcttcggtcaccacatccagcaccgccgcacacggaacaccggtggtggccagccagctcagacgcgccgcttcatc
ctgcagctcgttcagcgcaccgctcagatcggttttcacaaacagcaccggacgaccctgcgcgctcagacgaaacaccgccgcatc
agagcagccaatggtctgctgcgcccaatcatagccaaacagacgttccacccacgctgccgggctacccgcatgcaggccatcctg
ttcaatcatactcttcctttttcaatattattgaagcatttatcagggttattgtctcatgagcggatacatatttgaatgtatttagaaaaat
aaacaaataggggttccgcgcacatttccccgaaaagtgccacctaaattgtaagcggcggccgc

Supplemental data 6

Sequence of KmR-ori

ggccggccatttgcggccgctcttccgcttcctcgctcactgactcgctgcgctcggtcgttcggctgcggcgagcggtatcagctca
ctcaaaggcggtaatacggttatccacagaatcaggggataacgcaggaaagaacatgtgagcaaaaggccagcaaaaggccagg
aaccgtaaaaaggccgcgttgctggcgtttttccataggctccgcccccctgacgagcatcacaaaaatcgacgctcaagtcagaggt
ggcgaaacccgacaggactataaagataccaggcgtttccccctggaagctccctcgtgcgctctcctgttccgaccctgccgcttac
cggatacctgtccgcctttctcccttcgggaagcgtggcgctttctcatagctcacgctgtaggtatctcagttcggtgtaggtcgttcg
ctccaagctgggctgtgtgcacgaaccccccgttcagcccgaccgctgcgccttatccggtaactatcgtcttgagtccaacccggta
agacacgacttatcgccactggcagcagccactggtaacaggattagcagagcgaggtatgtaggcggtgctacagagttcttgaag
tggtggcctaactacggctacactagaagaacagtatttggtatctgcgctctgctgaagccagttaccttcggaaaaagagttggtag
ctcttgatccggcaaacaaaccaccgctggtagcggtggtttttttgtttgcaagcagcagattacgcgcagaaaaaaaggatctcaa
gaagatcctttgatcttttctacggggtctgacgctcagtggaacgaaaactcacgttaagggattttggtcatgagattatcaaaaagg
atcttcacctagatccttttaaattaaaaatgaagttttaaatcaatctaaagtatatatgagtaaacttggtctgacagttattagaaaaat
tcatccagcagacgataaaacgcaatacgctggctatccggtgccgcaatgccatacagcaccagaaaacgatccgcccattcgccg
cccagttcttccgcaatatcacgggtggccagcgcaatatcctgataacgatccgccacgcccagacggccgcaatcaataaagccg
ctaaaacggccattttccaccataatgttcggcaggcacgcatcaccatgggtcaccaccagatcttcgccatccggcatgctcgcttt
cagacgcgcaaacagctctgccggtgccaggccctgatgttcttcatccagatcatcctgatccaccaggcccgcttccatacgggta
cgcgcacgttcaatacgatgtttcgcctgatgatcaaacggacaggtcgccgggtccagggtatgcagacgacgcatggcatccgcc
ataatgctcactttttctgccggcgccagatggctagacagcagatcctgacccggcacttcgcccagcagcagccaatcacggcccg
cttcggtcaccacatccagcaccgccgcacacggaacaccggtggtggccagccagctcagacgcgccgcttcatcctgcagctcgt
tcagcgcaccgctcagatcggttttcacaaacagcaccggacgaccctgcgcgctcagacgaaacaccgccgcatcagagcagcca
atggtctgctgcgcccaatcatagccaaacagacgttccacccacgctgccgggctacccgcatgcaggccatcctgttcaatcatac
tcttcctttttcaatattattgaagcatttatcagggttattgtctcatgagcggatacatatttgaatgtatttagaaaaataaacaaatag
gggttccgcgcacatttccccgaaaagtgccacctaaattgtaagcgttaatattttgttaaaattcgcgttaaatttttgttaaatcagct
catttttaaccaataggccgaaatcggcaaaatccc
